# Aim18p and Aim46p are CHI-domain-containing mitochondrial hemoproteins in *Saccharomyces cerevisiae*

**DOI:** 10.1101/2022.11.15.516536

**Authors:** Jonathan M. Schmitz, John F. Wolters, Nathan H. Murray, Rachel M. Guerra, Craig A. Bingman, Chris Todd Hittinger, David J. Pagliarini

**Affiliations:** Department of Biochemistry, University of Wisconsin–Madison, Madison, WI, USA; Morgridge Institute for Research, Madison, WI, USA; Laboratory of Genetics, Genome Center of Wisconsin, Wisconsin Energy Institute, J.F. Crow Institute for the Study of Evolution, University of Wisconsin–Madison, Madison, WI, USA; DOE Great Lakes Bioenergy Research Center, University of Wisconsin–Madison, Madison, WI, USA; Department of Cell Biology and Physiology, Washington University School of Medicine, St. Louis, MO, USA; Department of Biochemistry and Molecular Biophysics, Washington University School of Medicine, St. Louis, MO, USA; Department of Genetics, Washington University School of Medicine, St. Louis, MO, USA

**Keywords:** Mitochondria, hemoprotein, chalcone isomerase, yeast, and uncharacterized protein

## Abstract

Chalcone isomerases (CHIs) have well-established roles in the biosynthesis of plant flavonoid metabolites. *Saccharomyces cerevisiae* possesses two predicted CHI-like proteins, Aim18p (encoded by YHR198C) and Aim46p (YHR199C), but it lacks other enzymes of the flavonoid pathway, suggesting that Aim18p and Aim46p employ the CHI fold for distinct purposes. Here, we demonstrate that Aim18p and Aim46p reside on the mitochondrial inner membrane and adopt CHI folds, but they lack select active site residues and possess an extra fungal-specific loop. Consistent with these differences, Aim18p and Aim46p lack chalcone isomerase activity and also the fatty acid-binding capabilities of other CHI-like proteins, but instead bind heme. We further show that diverse fungal homologs also bind heme and that Aim18p and Aim46p possess structural homology to a bacterial hemoprotein. Collectively, our work reveals a distinct function and cellular localization for two CHI-like proteins, introduces a new variation of a hemoprotein fold, and suggests that ancestral CHI-like proteins were hemoproteins.

## INTRODUCTION

Chalcone isomerases (CHIs) catalyze one of the essential steps in plant flavonoid biosynthesis (1–3). In the CHI reaction, bicyclic chalcones are isomerized into tricyclic flavanones (2, 4). This reaction is initiated by a catalytic arginine and supported through hydrogen bonding with other active site residues. Mutations to any of these residues reduce catalytic activity significantly (5).

Recently, CHIs were shown to belong to a larger family of CHI-domain-containing proteins that also contains a clade of non-catalytic fatty acid-binding proteins (6). Additionally, a recent study of reconstructed ancestral CHI-domain-containing proteins suggested that this protein family evolved from a non-catalytic ancestor (7), raising the possibility that there may be additional atypical CHI-domain-containing protein functions yet to be discovered.

Almost twenty years ago, Gensheimer and Musheigan discovered that the CHI-domain-containing protein family extends into fungi, slime molds, and γ-proteobacteria (8). Previous studies mention the presence of a fungal CHI-domain-containing protein family (6, 9–13) and speculate whether these proteins have CHI activity; however, to date, no functional characterization of the fungal CHI-domain-containing proteins has been reported.

*Saccharomyces cerevisiae* contains two paralogous CHI-domain-containing proteins, Aim18p (encoded by YHR198C) and Aim46p (encoded by YHR199C). More than a decade ago, a high-throughput screen in *S. cerevisiae* identified 100 genes whose absence disrupts the inheritance of mitochondrial DNA by daughter cells. Aim18p and Aim46p exceeded the threshold for this study, and therefore were assigned the standard name <u>A</u>ltered <u>I</u>nheritance of <u>M</u>itochondria (*AIM*). To our knowledge, no study has explicitly explored the molecular function of Aim18p and Aim46p in *S. cerevisiae*.

In this study, we find that Aim18p and Aim46p adopt the canonical CHI fold with an additional loop specific to fungi. We also show that Aim18p and Aim46p lack both chalcone isomerase activity and fatty acid-binding activity *in vitro*. Finally, we demonstrate that Aim18p and Aim46p are hemoproteins and that this heme-binding property is conserved widely among diverse fungi. Our findings expand the non-catalytic ancestor hypothesis by adding heme-binding to the list of functions of CHI-domain-containing proteins.

## RESULTS

### Aim18p and Aim46p are yeast proteins with sequence homology to plant CHIs

We sought to investigate whether *S. cerevisiae* possesses a flavonoid biosynthesis pathway by searching for homologs of plant flavonoid biosynthetic enzymes (14). We found distant homologs for multiple enzymes, but *S. cerevisiae* lacked obvious candidates for four pathway enzymes (Fig. 1A). Additionally, most of the *S. cerevisiae* homologs are known either to bind notably different substrates than their plant counterparts, or to perform entirely different biochemical reactions. For example, the *S. cerevisiae* homolog of chalcone synthase is Erg13p, an HMG-CoA synthase involved in ergosterol biosynthesis, not the biosynthesis of naringenin chalcone, the natural substrate for CHIs (Fig. 1A). Thus, while there are some distantly related, sequence-level homologs of some flavonoid biosynthesis genes (Table S1), this pathway most likely lacks functional conservation in *S. cerevisiae*.

**Figure 1.**
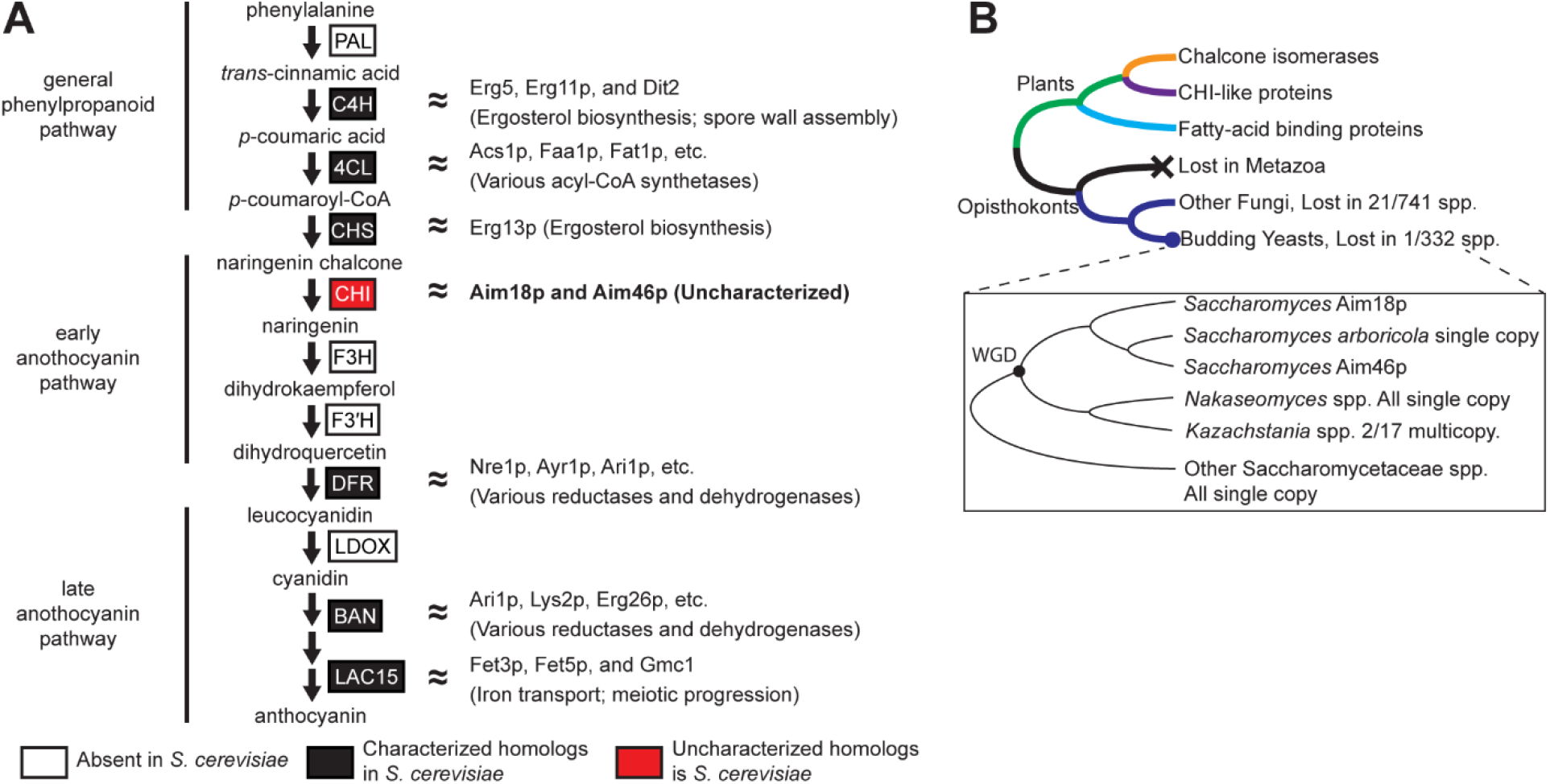
Aim18p and Aim46p are yeast proteins with sequence homology to plant CHIs. **A**. General flavonoid biosynthetic pathway with key enzymes and intermediates detailed. DELTA-BLAST searches for plant flavonoid biosynthetic pathway members were performed against the *Saccharomyces cerevisiae* genome, detailed in Table S1. White boxes indicate that *S. cerevisiae* homologs of plant flavonoid biosynthetic proteins are absent. Black boxes indicate that distant *S. cerevisiae* homologs of plant flavonoid biosynthetic proteins are characterized proteins; protein names and functions are noted at right. Red boxes indicate that *S. cerevisiae* homologs of plant flavonoid biosynthetic proteins are uncharacterized proteins. **B**. Simplified phylogenetic representation of CHI-domain-containing proteins across plants and fungi (Fig. S1A). Insert highlights species descended from the yeast whole genome duplication (WGD); note that no extant species retain the second copy from the WGD, and *AIM18* and *AIM46* are present in tandem in *Saccharomyces* spp., except for *S. arboricola*.

In contrast to the other flavonoid biosynthesis pathway homologs in *S. cerevisiae*, the CHI homologs Aim18p and Aim46p remain uncharacterized. Since *S. cerevisiae* does not have a functionally conserved chalcone synthase and likely lacks a functional flavonoid biosynthesis pathway, Aim18p and Aim46p probably do not perform the canonical CHI reaction.

Homologs of *S. cerevisiae* Aim18p and Aim46p are broadly distributed across the fungal kingdom. To interrogate the phylogenetic relationship between fungal and plant CHIs, we combined CHI plant protein sequences (7) and CHI homologs from over 1000 sequenced fungal genomes (15, 16), and we expanded the CHI-domain-containing protein family tree (6) (Fig. 1B; Fig. S1A). All fungal CHI-domain-containing proteins are an outgroup to the plant CHI-domain-containing proteins. Aim18p and Aim46p are encoded by paralogs that likely arose from a tandem gene duplication event specifically in the lineage leading to the genus *Saccharomyces*. The single copy present in the published *S. arboricola* genome likely resulted from a fusion of these paralogs, as evidenced by different phylogenetic affinities between its 5’ and 3’ ends (see Experimental Procedures). The proteins have significantly diverged in sequence, but both retain a clear CHI domain. Protein sequence alignments of AtCHI1, AtFAP1, Aim18p, and Aim46p demonstrate that Aim18p and Aim46p have sequence similarity in the more structured regions of previously crystallized CHI-domain-containing proteins (2, 6), especially the β3 [LXGXGXR] motif that houses the catalytic arginine (Fig. S1B).

### Aim18p and Aim46p are inner mitochondrial membrane proteins

Aim18p and Aim46p each contain N-terminal mitochondrial targeting sequences (Fig. 2A), as predicted by MitoFates (17), and an approximately 18 amino acid residue N-terminal transmembrane domain (Fig. 2A), as predicted by Phobius (18) (Fig. S2A). Crude mitochondrial purifications showed a strong mitochondrial enrichment for Aim18p-FLAG and Aim46p-FLAG compared to whole cell fractions (Fig. 2B), which is consistent with previous high-throughput studies that localized these proteins to mitochondria (19–22). The submitochondrial localization of Aim18p and Aim46p is ambiguous; early studies placed these proteins on the outer mitochondrial membrane (OMM) (23), while more recent studies suggest they reside on the inner mitochondrial membrane (IMM) (21, 22).

**Figure 2.**
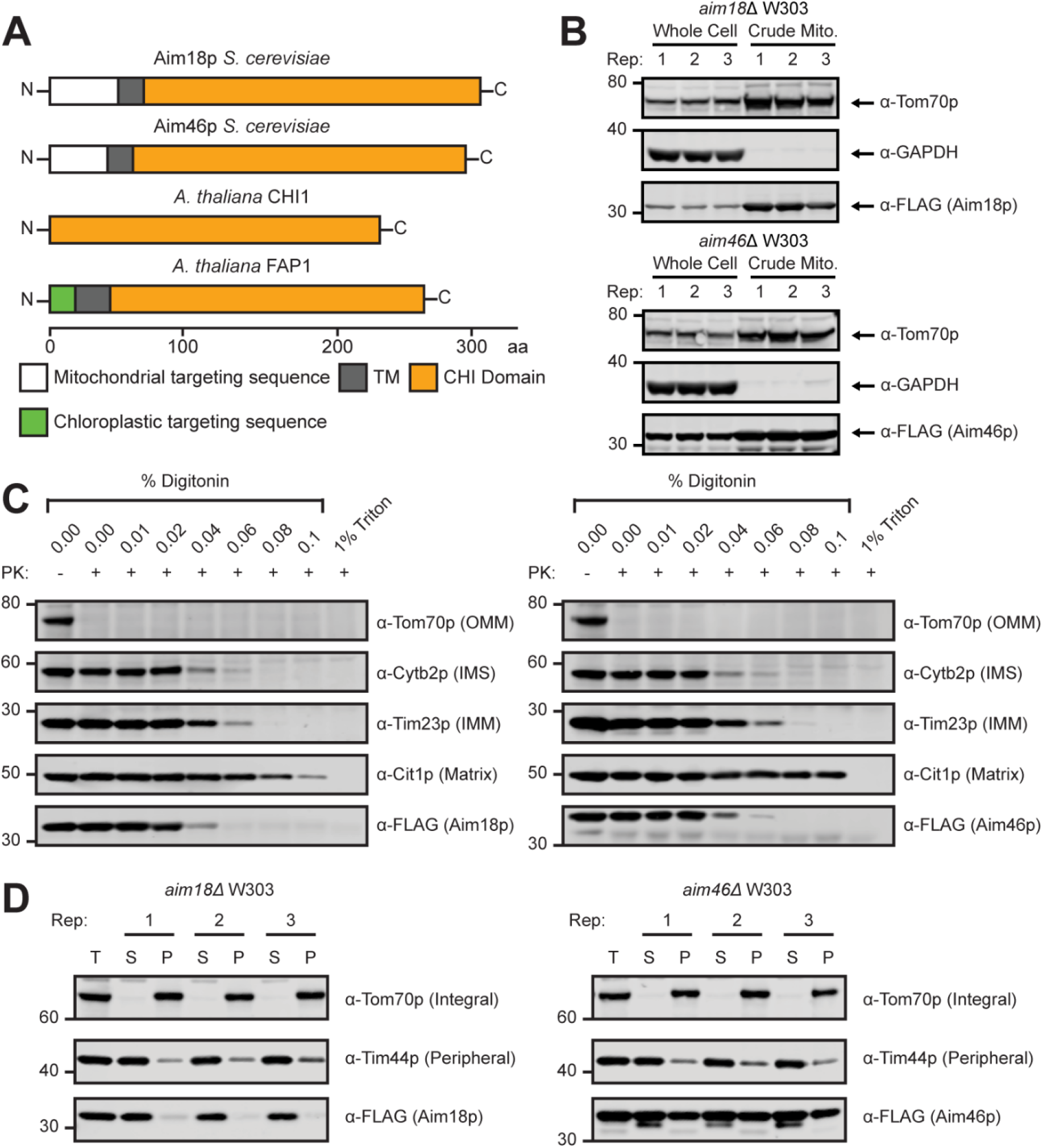
Aim18p and Aim46p are inner mitochondrial membrane proteins. **A**. Cartoon diagram of Aim18p, Aim46p, AtCHI1, and AtFAP1 protein sequences. Organellar targeting sequences, transmembrane domains, and CHI-domains are highlighted with their corresponding amino acid residue number. **B**. Western blot of Aim18p and Aim46p mitochondrial purifications compared to whole cells. Tom70p, mitochondrial; GAPDH, cytoplasmic; Aim18p-FLAG and Aim46p-FLAG, mitochondrial. *n* = 3 independent biological replicates. **C**. Proteinase K protection assay of yeast crude mitochondria expressing Aim18p-FLAG and Aim46p-FLAG. Yeast crude mitochondria were treated with 300 μg proteinase K in the presence of increasing concentrations of detergent. 1% triton was a control for full mitochondrial solubilization. OMM, outer mitochondrial membrane; IMS, intermembrane space; IMM, inner mitochondrial membrane. Representative blot from two independent experiments. **D**. Sodium carbonate extraction of yeast crude mitochondria expressing Aim18p-FLAG and Aim46p-FLAG. Yeast crude mitochondria were treated with 100 μL of 100 μM sodium bicarbonate for 30 minutes on ice. *n* = 3 independent technical replicates.

To clarify the localization of these proteins, we performed proteinase K protection assays on crude mitochondrial purifications expressing FLAG-tagged variants of Aim18p and Aim46p. Immunoblots for various mitochondrial markers and our FLAG-tagged target proteins suggest that Aim18p and Aim46p are localized to the IMM (Fig. 2C). A recent high-throughput study suggested that Aim18p and Aim46p might be peripherally associated with a membrane via their transmembrane domains (21). To determine whether Aim18p’s and Aim46p’s predicted hydrophobic regions behave as transmembrane domains or alternatively associate peripherally with the membrane in a manner similar to amphipathic helices, we performed sodium carbonate extractions on the same crude mitochondrial purifications used in Fig. 2C (24). Upon treatment with sodium bicarbonate, Aim18p was liberated into the soluble fraction, while a significant proportion of Aim46p remained in the insoluble fraction (Fig. 2D), suggesting that Aim46p’s transmembrane domain might be more strongly associated with the IMM.

*AIM18* and *AIM46* were previously implicated in mitochondrial inheritance in a high-throughput genetic screen (25), but we were unable to repeat this phenotype (Fig. S2B). Aim18p also exhibited correlated expression patterns with enzymes involved in coenzyme Q biosynthesis (26), but we did not observe altered coenzyme Q levels in *aim18* and/or *aim46* deletion mutants (Fig. S2C). We also did not observe any respiratory deficiency or growth phenotype in these strains (Fig. S2D), suggesting that Aim18p and Aim46p are not required for mitochondrial respiration.

### Aim18p and Aim46p adopt the CHI-fold but lack CHI-fold activities

In the absence of a growth phenotype, we employed biochemical techniques to explore the function of Aim18p and Aim46p. Full-length constructs of Aim18p and Aim46p were insoluble, but truncating the N-terminal MTS and TM for Aim18p and Aim46p (Aim18p-Nd70 and Aim46p-Nd62, respectively) dramatically increased solubility (Fig. 3A, 3B). These recombinant proteins were expressed in *E. coli* and purified to high concentrations (> 20 mg/mL in several instances) and were amenable to size exclusion chromatography. Both Aim18p and Aim46p had elution volumes similar to ~30 kDa proteins (See Experimental Methods; Fig. S3A), which suggests that they exist as monomers. We also generated point mutations of the CHI catalytic arginine—Aim18p-Nd70-R123A and Aim46p-Nd62-R107A—to serve as experimental controls. Each protein exhibited robust thermal melt curves, indicating stably folded protein (Fig. S3B). Wild-type Aim46-Nd62 readily formed high-quality crystals, while only the arginine mutant of Aim18-Nd70 formed crystals, presumably due to the increased stability (Fig. S3B).

**Figure 3.**
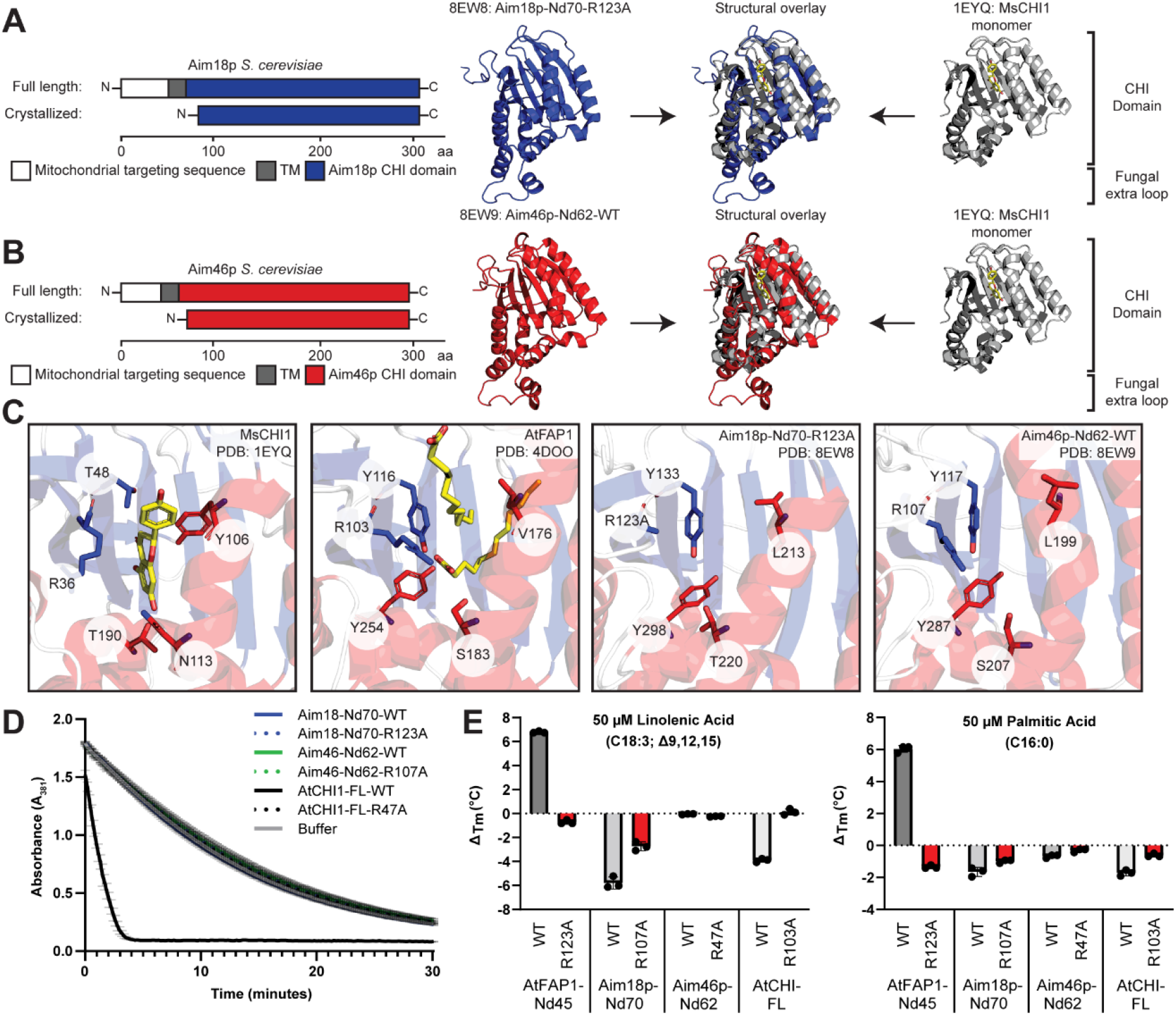
Aim18p and Aim46p adopt the CHI-fold but lack CHI-fold activities. **A**. Diagram of crystallized Aim18p (Blue) compared to the full-length protein from *S. cerevisiae* and structural overlay with CHI from *M. sativa* (Gray, PDB: 1YEQ)*. M. sativa* CHI naringenin chalcone ligand highlighted in yellow. **B**. Cartoon of crystallized Aim46p (Red) compared to the full-length protein from *S. cerevisiae* and structural overlay with CHI from *M. sativa* (Gray, PDB: 1YEQ)*. M. sativa* CHI naringenin chalcone ligand highlighted in yellow. **C**. Active site comparisons between MsCHI1, AtFAP1, Aim18p, and Aim46p. Bound ligands for MsCHI1 (naringenin) and AtFAP1 (laurate) highlighted in yellow. Alpha helices colored in red, beta sheets colored in blue, and linker regions colored in gray. **D**. Chalcone isomerase assay with 10 nM CHI-domain-containing proteins and 200 μM naringenin chalcone. Conversion of naringenin chalcone to naringenin measured by A381 readings taken every 10 seconds for 30 minutes (mean ± SD, *n* = 3 independent technical replicates). **E.** Thermal shift response of 5 μM CHI-family proteins in response to a 1-hour incubation with 50 μM fatty acids compared to a DMSO control (mean ± SD, *n* = 3 independent technical replicates).

We solved the crystal structures for Aim18p-Nd70-R123A (PDB: 8EW8) and Aim46p-Nd62-WT (PDB: 8EW9) to resolutions of 2.15 Å and 2.0 Å, respectively (Table S3). Both Aim18p-Nd70-R123A and Aim46p-Nd62-WT adopt the CHI fold, as determined by similarity to CHI1 from *Medicago sativa* (MsCHI1) (Fig. 3A-B; PDB: 1EYQ (2)). Almost the entirety of the core 7-stranded antiparallel β-sheet is shared between these proteins with small deviations in the more solvent-accessible α-helices (Fig. 3A-B). Of note is a region of the Aim18p-Nd70 and Aim46-Nd62 structures that lies in between plant CHI α1 and α2. This region contains an extra loop that is absent in plant CHI-domain-containing proteins (Fig. 3A-B; Fig. S3D) but is conserved across multiple ascomycete Aim18p and Aim46p homologs (Fig. S3E).

Nearly all of the CHI catalytic residues are substituted non-conservatively in Aim18p and Aim46p except for the catalytic arginine (Fig. 3C), which is invariably conserved in fungal homologs of Aim18p and Aim46p (Fig. S3C). A single non-conservative mutation in these residues is sufficient to nearly abolish catalytic activity (5), so the presence of several substitutions suggests that Aim18p and Aim46p likely lack canonical CHI enzymatic activity. In contrast, Aim18p and Aim46p share 4/5 active site residues with the fatty acid-binding CHI-like protein, FAP1 from *Arabidopsis thaliana* (AtFAP1) (PDB: 4DOO (6); Fig. 3C; Fig. S3C).

To test whether Aim18p and Aim46p possess either CHI or fatty acid-binding activities, we first designed protein expression constructs for the control proteins AtCHI1 and AtFAP1-Nd45 (full-length AtFAP1 did not express, much like the fungal CHI-domain-containing proteins, which require an N-terminal truncation for purification). We performed the standard CHI assay on wild-type and catalytic arginine mutant constructs of Aim18p-Nd70, Aim46p-Nd62, and AtCHI1 (Fig. 3D; Fig. S4A). Only AtCHI-WT rapidly converted naringenin chalcone to naringenin, supporting the hypothesis that Aim18p and Aim46p lack CHI catalytic activity. We also performed differential scanning fluorimetry to assess changes in thermal stability on wild-type and catalytic arginine mutant constructs of Aim18p-Nd70, Aim46p-Nd62, AtCHI1, and AtFAP1-Nd45 as a proxy for lipid binding (6) (Fig. 3E; Fig. S4A). The positive control AtFAP1-Nd45-WT was the only protein stabilized by the addition of linolenic acid and palmitic acid, while the rest of the proteins were either not stabilized or were destabilized, a phenomenon also observed with AtCHI1 (6). Taken together, while Aim18p and Aim46p adopt the canonical CHI fold, the lack of CHI catalytic activity and absence of fatty acid-biding activity suggest that fungal CHI-domain-containing proteins likely have a different biological activity from plant CHI-domain-containing proteins.

### Aim18p and Aim46p are hemoproteins

Our purified recombinant Aim18p-Nd70 and Aim46p-Nd62 from *E. coli* exhibited a golden color, suggesting a bound ligand (Fig. 4A; Fig. S4A). Interestingly, catalytic arginine mutants lacked this golden color. Spectroscopy revealed that the colored fungal CHI-domain-containing proteins had a distinct peak around 420 nm (Fig. 4B; Fig. S4D-G), which likely represents the signature Soret peak of hemoproteins (27).

**Figure 4.**
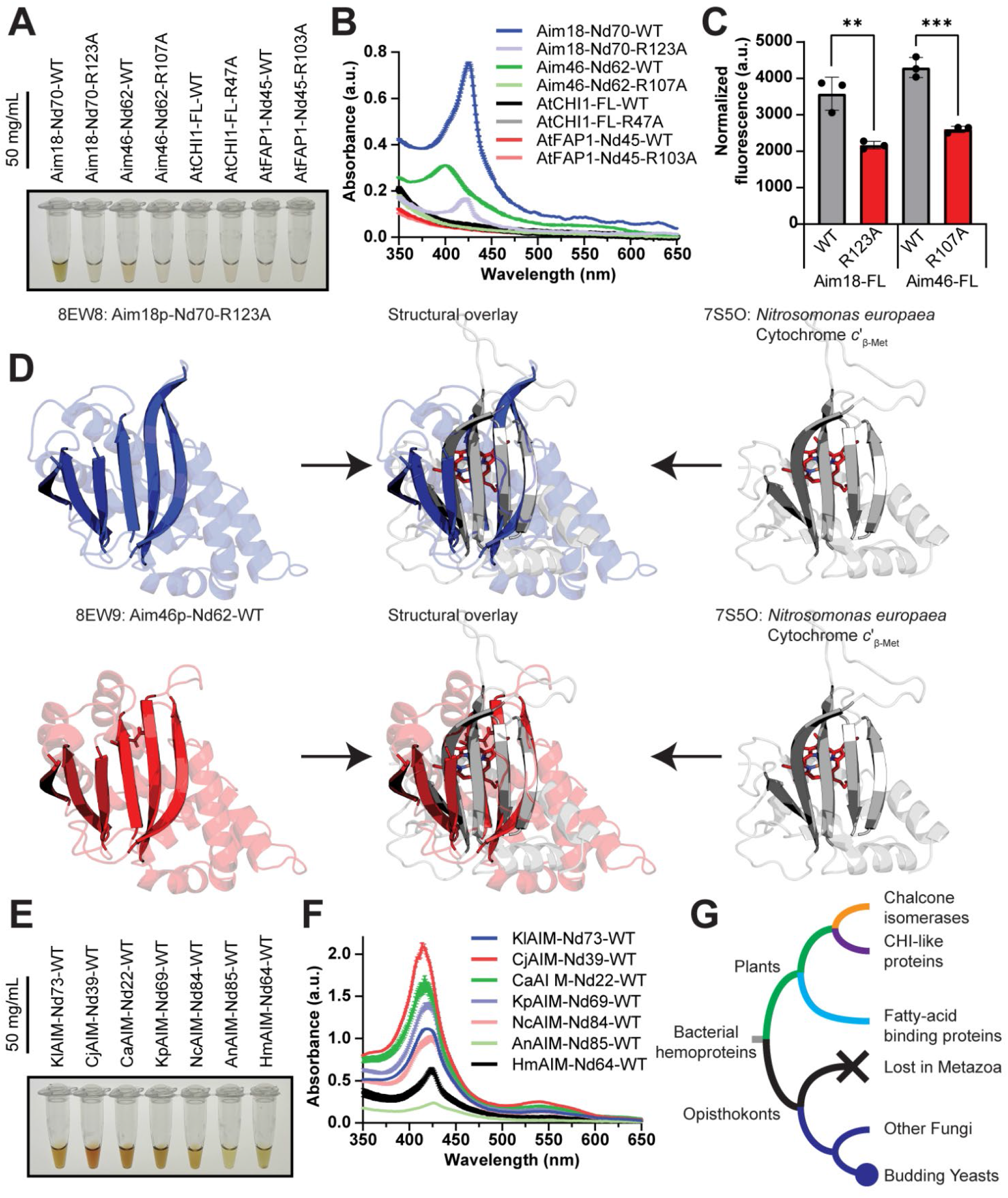
Aim18p and Aim46p are hemoproteins. **A**. Image of 50-mg/mL recombinant Aim18p-Nd70 and Aim46p-Nd62 variants co-purifying with a golden color compared to AtCHI1 and AtFAP1. **B**. UV-Vis spectroscopy of proteins in Fig. 4A highlighting the presence of a “Soret” peak at ~400 nm, a characteristic of hemoproteins. Mean ± SD, *n* = 3 independent technical replicate measurements of the same sample. **C**. Fluorometric oxalic acid heme determination assay with immunoprecipitations of FLAG-tagged Aim18p and Aim46p expressed in *S. cerevisiae* (**p = 0.0062 Aim18-FL-WT vs Aim18-FL-R123A, ***p = 0.0002 Aim46p WT vs Aim46p R107A; mean ± SD, *n* = 3 independent technical replicates). Average heme content measurements were normalized via densitometry by protein abundance (Fig. S4C). Significance was calculated by an unpaired, two-tailed Student’s t-test. **D**. Structural overlay of Aim18p (Blue) and Aim46p (Red) with cytochrome c’_β-Met_ from the bacterium *N. europaea* (Gray, PDB: 7S5O). Alpha helices made slightly transparent to highlight homology of core antiparallel beta sheet for both aligned proteins. **E**. Image of recombinant fungal AIM homologs co-purifying with golden color. Recombinant fungal AIM homologs are named based on their species of origin (KlAIM = *K. lactis*; CjAIM = *Cy. jadinii*; CaAIM = *C. albicans*; KpAIM = *Ko. pastoris*; NcAIM = *N. crassa*; AnAIM = *A. nidulans*; HmAIM = *H. marmoreus*). **F**. UV-Vis spectroscopy of proteins in Fig. 4E highlighting the presence of a “Soret” peak at ~400 nm, a characteristic of hemoproteins. Mean ± SD, *n* = 3 independent technical replicate measurements of the same sample. Recombinant fungal AIM homologs are named based on their species of origin (KlAIM = *K. lactis*; CjAIM = *Cy. jadinii*; CaAIM = *C. albicans*; KpAIM = *Ko. pastoris*; NcAIM = *N. crassa*; AnAIM = *A. nidulans*; HmAIM = *H. marmoreus*). **G**. Proposed phylogenetic tree of CHI-domain-containing proteins with bacterial hemoproteins as the outgroup.

We considered whether heme binding might be an artifact of expressing recombinant fungal CHI-domain-containing proteins in *E. coli*. To assuage this concern, we generated centromeric yeast expression plasmids encoding FLAG-tagged, full-length Aim18p and Aim46p to test heme binding in their native organism, *S. cerevisiae*. Due to the lower yield of protein from these immunoprecipitations, we confirmed the presence of heme through an orthogonal approach. We used a fluorescent heme detection assay that uses heated oxalic acid to liberate the heme molecule of its central iron metal, generating a fluorescent protoporphyrin molecule. This experiment revealed that Aim18p and Aim46p co-immunoprecipitated with bound heme and that mutating the catalytic arginine significantly reduced heme binding (Fig. 4C; Fig. S4C). Despite multiple attempts, we were unable to generate heme-bound crystal structures for either Aim18p or Aim46p. Additionally, the structure of Aim46p-Nd62-WT contained a small molecule resembling α-ketoglutarate in the active site, but we were unable to confirm its identity via mass spectrometry.

The tertiary structure of the CHI domain is classified as a mixed α+β fold. To explore whether other hemoproteins possess an α+β fold similar to Aim18p and Aim46p, we searched our structures against the Dali server (28). A heme-c-binding protein, cytochrome *c*’_β-Met_ from the ammonia-oxidizing bacterium *Nitrosomonas europaea* (PDB: 7S5O; (29)), was among the top hits outside of the plant CHI-domain-containing proteins. Alignments of the Aim18-Nd70-R123A and Aim46-Nd62-WT structures with the bacterial protein revealed that, while the α-helices differed substantially, the core CHI antiparallel β-sheet was structurally homologous to the core antiparallel β-sheet of cytochrome *c*’_β-Met_ (Fig. 4D).

The presence of a bacterial hemoprotein with structural similarities to the CHI fold suggests the possibility that the ancestral CHI protein was a hemoprotein. To further determine the phylogenetic breadth of heme-binding, we leveraged the rich dataset of fungal CHI-domain-containing proteins (Fig. S1A) to find extant fungal CHI-domain-containing proteins representing several clades of fungi (Fig. S3C), including at least one representative from the phylum Basidiomycota and all three subphyla of the phylum Ascomycota. With these sequences, we designed N-terminally truncated (18) protein expression constructs for these fungal CHI-domain-containing proteins. We observed that all successfully purified recombinant fungal CHI-domain-containing proteins co-purified with heme (Fig. 4E-F; Fig. S4B; Fig. S4D-G). Collectively, these data suggest that the ancestral fungal CHI-domain-containing protein was a hemoprotein, and that the entire CHI-domain-containing protein family might share a common ancestor with a family of bacterial hemoproteins (Fig. 4G).

## DISCUSSION

CHI-domain-containing proteins (CHIs) comprise a large family of proteins that feature a common fold but wield multiple functions. This family derives its name from enzymes that catalyze the intermolecular cyclization of chalcones to flavanones (Type I and Type II CHIs; (2, 30)), but this family also includes fatty-acid-binding proteins (Type III CHIs; (6)) and non-catalytic proteins that are thought to increase flavonoid biosynthesis (Type IV CHIs; (31, 32)). In this study, we expand the functional repertoire of the CHI-domain-containing protein family by revealing that the fungal clade of CHIs are hemoproteins that lack canonical CHI activity. Moreover, given that gene duplications are less widespread among budding yeasts than among the CHI-domain containing proteins seen in plants, it is likely that these fungal sequences are more conserved in function and may be more functionally similar to the ancestral protein (Fig. S1A).

Despite these advances, without an *in vivo* phenotype or heme-bound crystal structure, it is difficult to ascertain how Aim18p and Aim46p employ their CHI fold and heme-binding capabilities in mitochondria. Hemoproteins have multiple functions, including, but not limited to, ligand delivery (e.g., oxygen via hemoglobin), signal transduction (e.g., nitric oxide binding by soluble guanylyl cyclases), electron transport, and catalysis (e.g., peroxidases and monooxygenases) (33, 34).

The CHI fold is classified as an α+β fold, which comprises a mostly antiparallel beta-sheet decorated with alpha helices (35). α+β hemoproteins (including Aim18p and Aim46p) comprise approximately 10% of all known hemoprotein folds (36, 37) and often leverage their heme for environmental sensing and heme transport functions. These include the Per-Arnt-Sim (PAS)-domain-containing hemoproteins (38, 39), such as the oxygen sensors Dos and FixL (40, 41). Other examples of α+β hemoproteins are the heme acquisition protein HasA in bacteria (42, 43) and the SOUL family of hemoproteins, which have proposed biological functions ranging from regulating cell fate to heme transport (44). There are exceptions to this theme, including the multifunctional α+β ferredoxin proteins (45, 46), but the majority of α+β hemoproteins are gas sensors or heme transporters. While we do not know the exact function of Aim18p and Aim46p, it is reasonable to speculate that these proteins might regulate mitochondrial function by utilizing heme in a similar manner.

We observed that the heme binding of recombinant Aim46p, when purified from *E. coli*, was more variable across preps (Fig. S4D, F). This suggests that heme binding might be sensitive to environmental fluctuations and perhaps that Aim18p and/or Aim46p might serve as heme-sensing proteins, although this remains speculation. Although our data show that the conserved CHI arginine does influence heme binding (Fig. 4A-C; Fig. S4D, F) in Aim18p and Aim46p, it is most likely not the only heme-coordinating residue; future experiments will be necessary to fully understand the coordination environment between heme and the CHI domain.

Our investigation of Aim18p and Aim46p is part of the grand post-genomic era effort to define functions for all proteins. (47, 48). Modern structural biology approaches and the advent of powerful structural prediction algorithms doubtlessly are advancing us toward this goal (49–51). However, our work serves as a reminder that biochemical functions are often not immediately evident from structure. We demonstrate that the CHI-domain-containing hemoproteins in fungi do not behave like their plant relatives, even though they have high sequence and structural homology. Protein folds conferring multiple functions is a well-established biological phenomenon (52); for example, thousands of proteins contain a Rossman-like fold, but their substrates, cofactors, and functions can differ significantly (53). Other examples of many include the α+β barrel hemoproteins, which share a ferredoxin-like fold, but have diverse functions (45, 46), as well as cytochrome P460 (an enzyme) and cytochrome c’_β_ (a gas-binding protein), which adopt a similar fold but bind different substrates (54).

To our knowledge, this is the first published biochemical study of a CHI-domain-containing protein in the fungal kingdom. Our analyses assign a distinct heme-binding function to the previously uncharacterized proteins Aim18p and Aim46p in *S. cerevisiae*, as well as to a diverse array of fungal homologs, expanding previous observations that the CHI domain is functionally plastic (6, 7, 31). We also suggest, through structural homology prediction, that the CHI domain might be related to bacterial cytochrome P460s and suggest that the ancestral CHI-like proteins may have relied on heme-binding for its function. Collectively, our study provides a foundation for further work to define the evolution and specific *in vivo* roles of Aim18p and Aim46p.

## EXPERIMENTAL PROCEDURES

### Evolutionary Analysis

Flavonoid biosynthesis proteins from *A. thaliana* were used to search for homologs in *S. cerevisiae* using DELTA-BLAST with an e-value < 1e-15. To search for CHI homologs in yeasts, a HMMER profile was constructed using *S. cerevisiae* Aim18p and Aim46p and used to conduct a HMMER search 332 annotated budding yeast genomes (15) to identify hits with score >50 and e-value <0.001. To search for CHI homologs in more divergent fungi, tBLASTn was used to identify hits in over 700 unannotated Ascomycota genomes (16) with e-value <1e-6. The full plant and fungal CHI homolog phylogeny was built by adding the fungal sequences to a structural alignment of plant CHI homolog sequences (7) using the—add function of MAFFT (55) using default settings. The resulting alignment was truncated to the regions matching the initial structural alignment that included the CHI domain and filtered to sites with less than 50% gaps using trimAl (56). A phylogenetic tree was constructed using fasttree (57) with the settings (d -lg -gamma -spr 4 -slownni -mlacc 2), midpoint rooted, and edited to collapse clades using iTOL(58). The absence of any CHI homologs in metazoan lineages was determined by repeated search attempts using both blastp and DELTA-BLAST at permissive e-value thresholds restricted to relevant taxonomic groups. The determination that the single CHI homolog in the published genome of *S. arboricola* was due to a fusion of paralogs was made by aligning the CDS of all hits from *Saccharomyces* species via MAFFT using default settings (55) and looking for signatures of recombination between *AIM18* and *AIM46* homologs in the *S. arboricola* gene using RDP4 (59). A single recombination event was found in this sequence (all tests of recombination were significant at p<1e-13), which supports a fusion with *AIM46* providing roughly the 5’ 60% and *AIM18* the final 3’ 40% of the sequence.

A subset of protein sequences of plant and fungal CHIs were aligned via MAFFT using default settings (55). Protein sequence alignments were visualized using Jalview 2.11.2.5 and colored by sequence conservation (60). Mid-point rooted phylogenetic trees of plant and fungal CHIs were generated through the Phylogeny.fr “one click” mode workflow (http://phylogeny.lirmm.fr/) using default settings (61–65). Alignments and phylogenetic trees were exported as svg files and annotated in Adobe Illustrator 26.5.

### Yeast Strains and Cultures

All yeast strains used in this study were (or were derived from) wild-type *Saccharomyces cerevisiae* haploid W303-1A (*MAT***a**, *his3 leu2 met15 trp1 ura3*). Yeast single and double deletions of *aim18* and/or *aim46* were generated by replacing the ORFs with the *HIS3MX6* cassette via homologous recombination, as previously described (66) (Table S2). Homologous recombination donor DNA was ~5 μg *HIS3MX6* cassette amplified with 40 bp homology upstream and downstream of the corresponding ORF. Successful insertion of the knockout cassette was confirmed by a PCR-based genotyping assay.

Unless otherwise described, yeast strains were cultured in synthetic complete (SC) (or synthetic complete minus uracil (Ura^-^), for plasmid-based experiments) medium with the indicated carbon source. SC medium contained drop-out mix complete (without yeast nitrogen base, US Biologicals: D9515), yeast nitrogen base (without amino acids, carbohydrate, or ammonium sulfate, US Biologicals: Y2030), 5 g/L ammonium sulfate, and the indicated carbon source (w/v). SC Ura^-^ medium contained drop-out mix synthetic minus uracil (without yeast nitrogen base, US Biologicals: D9535), yeast nitrogen base (without amino acids, carbohydrate, or ammonium sulfate, US Biologicals: Y2030), 5 g/L ammonium sulfate, and the indicated carbon source (w/v). All media pH values were adjusted to 6.0 and sterilized via vacuum filtration (0.22 μM).

### Plasmid Cloning

Yeast expression plasmids were cloned via standard restriction enzyme and ligation methods. ORF-specific inserts for *AIM18* and *AIM46* (and their respective arginine mutants) with 20 bp homology to p416-GPD (Betty Craig Lab) were amplified from W303 yeast genomic using primers specific to each ORF (Table S2). PCR-amplified ORF inserts were ligated into *Xba*I- and *Sal*I-HF-digested p416 GPD and then then transformed into *E. coli* 10 G chemically competent cells (Lucigen). Successful transformants were miniprepped and confirmed via Sanger sequencing (UW–Madison Biotechnology DNA sequencing core facility). For experiments, yeast strains were transformed with ~300 ng plasmid via the LiAc/SS-DNA/PEG method (Geitz and Woods, 2002).

*E. coli* protein expression plasmids were cloned via standard Gibson assembly. ORF inserts with 40 bp homology to pET-21a(+) were amplified from W303 yeast genomic DNA (for *AIM18* and *AIM46*) or gBlocks (for *AtCHI1, AtFAP1*, and fungal *AIM* homologs) using 60 bp primers specific to each ORF (Table S2). gBlocks for fungal *AIM* homologs included *AIM* ORFs from: *Kluyveromyces lactis, Cyberlindnera jadinii, Candida albicans, Komagataella pastoris, Yarrowia lipolytica, Lipomyces starkeyi, Neurospora crassa*, *Aspergillus niger, Schizosaccharomyces pombe*, and *Hypsizygus marmoreus* (Table S2; Supporting Documents S1.zip). PCR-amplified ORF inserts were Gibson assembled into *Nco*I-HF- and *Xho*I-digested pET-21a(+) (Addgene) and then transformed into *E. coli* 10 G chemically competent cells (Lucigen). Successful transformants were miniprepped and confirmed via Sanger sequencing (UW–Madison Biotechnology DNA sequencing core facility).

### Site-Directed Mutagenesis

Site-directed mutagenesis (SDM) reactions to generate catalytic arginine point mutants for *AIM18* and *AIM46* were performed according to the Q5® Site-Directed Mutagenesis Kit (New England Biolabs, Table S2). SDM reactions were transformed into *E. coli* 10 G chemically competent cells (Lucigen). Successful transformants were miniprepped and confirmed via Sanger sequencing (UW–Madison Biotechnology DNA Sequencing Core Facility). Arginine point mutants for *AtCHI1* and *AtFAP1* were ordered as separate gBlocks.

### Yeast Drop Assay

Single colonies of *S. cerevisiae* W303-1A were used to inoculate 3-mL starter cultures of SC +2% glucose (w/v) medium, which were incubated overnight spinning on culture tube wheel (20 h, 30 °C, max speed). Yeast from these starter cultures were serially diluted (10^5^, 10^4^, 10^3^, 10^2^, and 10^1^) in water. 4 μL of each serial dilution were dropped onto solid agar SC plates containing either 2% glucose (w/v), 3% glycerol (w/v), or 2% EtOH (v/v) and incubated (2-4 days, 30 °C).

### Yeast Petite Frequency Assay

*S. cerevisiae* W303-1A were streaked onto non-fermentable YPEG plates to eliminate petites (1% yeast extract [w/v], 2% peptone [w/v], 3% EtOH [w/v], 3% glycerol [w/v], and 2% agar [w/v]). Single colonies of petite-free *S. cerevisiae* W303-1A were used to inoculate 3-mL fermentation starter cultures of SC +2% glucose (w/v) medium, which were incubated overnight spinning on a culture tube wheel to remove selection against petite formation (30 °C, maximum speed). Yeast cells from these starter cultures were then plated for single colonies on YPDG plates (1% yeast extract [w/v], 2% peptone [w/v], 0.1% glucose [w/v], 3% glycerol [w/v], and 2% agar [w/v]) (2-3 days, 30 °C). Petite colonies were identified and counted manually, and the resulting petite frequencies were recorded in Microsoft Excel and analyzed in Prism 9.2.0 (GraphPad by Dotmatics).

### Crude Mitochondria Purification

Single colonies of *S. cerevisiae* W303-1A were used to inoculate 3-mL starter cultures of either SC +2% glucose (w/v) or Ura^-^ +2% glucose (w/v) (for plasmid-transformed strains) medium, which were incubated overnight spinning on a culture tube wheel (20-24 h, 30 °C, max speed). Starter cultures of yeast strains were used to inoculate 500-mL respiratory cultures of either SC +3% glycerol (w/v) +0.1% glucose (w/v) or Ura^-^ +3% glycerol (w/v) +0.1% glucose (w/v) (for plasmid-transformed strains) medium with 2.5 × 10^7^ cells, which were incubated until past the diauxic shift (25 h, 30 °C, 225 rpm). For whole-cell analysis, 1 × 10^8^ cells were collected, harvested by centrifugation (21,130 × *g*, 3 min, room temperature [RT]), flash frozen in N_2_(l), and stored at −80 °C. The remaining culture was harvested by centrifugation (4,000 × *g*, 10 min, RT), washed with 25 mL ddH_2_O, and then weighed (~2 g). Yeast crude mitochondria were then isolated as previously described (67). Isolated yeast crude mitochondria were gently resuspended (using a wide-bore pipette tip) in SEM buffer (250 mM sucrose, 1 mM EDTA, 10 mM MOPS-KOH, pH 7.2, 4 °C). Total protein concentration for whole-cell and isolated yeast crude mitochondria samples were quantified for total protein using the Pierce™ BCA Protein Assay Kit (Thermo). Isolated yeast crude mitochondria were aliquoted as 40 μg aliquots, flash frozen in N_2_(l), and stored at −80 °C.

### Yeast FLAG Immunoprecipitation

Single colonies of *S. cerevisiae* W303-1A expressing full-length Aim18p or Aim46p under control of the glyceraldehyde-3-phosphate dehydrogenase gene (THD3) promoter were used to inoculate 3-mL starter cultures of Ura^-^ +2% glucose (w/v) medium, which were incubated overnight spinning on a culture tube wheel (24 h, 30 °C, maximum speed). Starter cultures of yeast strains were used to inoculate 100-mL respiratory cultures of Ura^-^ +3% glycerol (w/v) +0.1% glucose (w/v) (for plasmid-transformed strains) medium supplemented with 500 μM ammonium iron (II) sulfate hexahydrate ([Argos Organics: 201370250], as iron is limiting in synthetic media) with 5 × 10^6^ cells, which were incubated until past the diauxic shift (34 h, 30 °C, 225 rpm). Yeast cultures were harvested by centrifugation (4,000 x *g*, 10 min, RT), washed with 25 mL ddH_2_O, transferred to a 2 mL screw-cap Eppendorf tube, and pinned via centrifugation (21,130 x *g*, 2 min, RT). Yeast pellets were resuspended in 400 μL FLAG lysis buffer (20 mM HEPES, pH 7.4, 100 mM NaCl, 10% [w/v] glycerol, 3% [w/v] digitonin [Sigma: D141], 1 mM DTT, 1X PI-1 [10 μM benzamidine, 1 μg/mL 1,10-phenanthroline], 1X PI-2 [0.5 μg/mL each of pepstatin A, chymostatin, antipain, leupeptin, aprotinin]), and 200 μL glass beads. Yeast pellet resuspensions were lysed via bead beating on a Disruptor Genie® (Zymo Research) cell disruption device (3X, 2 min, with 2 min rests in between bead beatings). Yeast lysates were clarified via centrifugation (21,130 x *g*, 15 min, 4 °C) and transferred to a new tube. Equal volumes of clarified yeast lysates (400 μL) were incubated with 20 μL Anti-FLAG® M2 Magnetic Beads (Millipore: M8823). Magnetic beads containing FLAG-tagged protein were washed four times with FLAG wash buffer (20 mM HEPES, pH 7.4, 100 mM NaCl, 0.05% [w/v] digitonin [Sigma: D141], and 10% [w/v] glycerol) and eluted with FLAG elution buffer (20 mM HEPES, pH 7.4, 100 mM NaCl, 0.05% [w/v] digitonin [Sigma: D141], 10% [w/v] glycerol, and 0.4 mg/mL, and 1x FLAG-peptide [Sigma: F3290]) (30 min, with moderate agitation, RT). 0.5% of each elution was analyzed via western blot as described below to check for equal protein expression and immunoprecipitation. For heme measurement assays, protein abundances were normalized to the first replicate of wild-type W303-1A yeast via densitometry with the ImageJ 1.53c software ((68), NIH).

### Proteinase K Protection Assay

Proteinase K protection assays were performed as previously described (Sato and Mihara, 2010). 40 μg yeast crude mitochondria were incubated with digitonin (0%, 0.01%, 0.02%, 0.04%, 0.06%, 0.08%, and 0.1% [w/v]) and 1% triton (w/v) in the presence of 300 μg/mL proteinase K (40 μL reaction size, 15 min, RT). Proteinase K was then inhibited with the addition of 7 mM PMSF. Each reaction was precipitated with 25% TCA (20 min, on ice) and resuspended in in 40 μL 1X LDS sample buffer containing 4% β-mercaptoethanol (BME). 10 μL of each reaction was analyzed via western blot as described below.

### Sodium Carbonate Extraction Assay

Sodium carbonate extraction assays were performed as previously described (24). 40 μg yeast crude mitochondria were incubated with 100 μL of 100 mM sodium bicarbonate (Fisher Scientific: S263, 30 min, on ice). Soluble and insoluble fractions were isolated via centrifugation (90,000 x *g*, 30 min, 4 °C) in an Optima™ MAX-OP (Beckman Coulter) centrifuge using the TLA-100.3 rotor. Pellets (insoluble fraction) and supernatants (soluble fraction) were precipitated with 25% TCA (20 min, on ice) and resuspended in in 40 μL 1X LDS sample buffer containing 4% β-mercaptoethanol (BME). 10 μL of each reaction was analyzed via western blot as described below.

### Western Blotting

Protein samples in 1X LDS sample buffer containing 4% β-mercaptoethanol (BME) were separated by size on 4-12% Novex NuPage Bis-Tris SPS-PAGE (Invitrogen) gels (2 h, 100 V). Gels were transferred to a Immobilon®-FL PVDF Membranes (Millipore) using the Mini Blot Module western blotting system (Invitrogen) for 1 h in transfer buffer (192 mM glycine, 25 mM Tris, 20% methanol [v/v], 4 °C). Membranes were dried for 15 min in the fume hood, rehydrated with a 15 second methanol wash, and submerged in TBST (500 mM NaCl, 20 mM Tris pH 7.4, 0.05% Tween 20 [v/v]). Membranes were blocked in 5% non-fat dry milk (NFDM) in TBST for 1 h on a rocker at RT. Membranes were incubated with primary antibodies diluted (Table S2) in 1% NFDM in TBST with 0.02% (w/v) sodium azide overnight on a rocker at 4 °C. After the primary incubation, membranes were washed in TBST (3X, 5 min, RT) on a rocker. Membranes were incubated on a rocker with fluorophore-conjugated secondary antibodies diluted (Table S2) in 5% NFDM in TBST (1 h, RT). After the secondary incubation, membranes were washed in TBST (3X, 5 min, RT) on a rocker and imaged on LI-COR Odyssey CLx using the Image Studio v5.2 software. Special care was made to ensure that membranes were exposed to the light as little as possible to protect the fluorescent signal of the secondary antibody.

### Antibodies

Proteins of interest were detected with primary antibodies to Tom70p (69) (a gift from Nora Vögtle, 1:500), GAPDH (ThermoFisher, MA5-15738, 1:4000), FLAG (Proteintech, 66008-3-lg, 1:5000), Cytb2 (70) (a gift from Betty Craig, 1:1000), Tim23 (71) (a gift from Betty Craig, 1:5000), Tim44 (72) (a gift from Betty Craig, 1:1000), and Cit1 (73) (Biomatik, 1:4000). Fluorescent conjugated secondary antibodies included α-mouse 800 (LICOR, 926-32210, 1:15000) and α-rabbit 800 (LICOR, 926-32211, 1:15000). Detailed information about antibodies is in Table S2.

### Purification of Recombinant Proteins

pET-21a(+) constructs containing DNA encoding 8X-His-tagged CHI-domain-containing proteins were transformed into BL21-CodonPlus (DE3)-RIPL Competent Cells (Agilent) for recombinant protein expression. Proteins were gently expressed overnight by autoinduction (74) (37 °C, 3 h; 20 °C, 20 h, 225 rpm). Cells were harvested via centrifugation (4000 x *g*; 30 min, RT), flash frozen in N_2_(l), and stored at −80 °C. For purification, pellets expressing recombinant protein were thawed on ice for at least an hour with lysis buffer (400 mM NaCl, 160 mM HEPES (pH 7.5), 5 mM BME, 0.25 mM PMSF, and 1 mg/mL lysozyme). Thawed pellets were resuspended via vortexing until the solution was homogenous. Resuspended pellets were lysed via sonication (3X, 70% amplitude, 20 s, with 60 s rests) and clarified via centrifugation (15000 x *g*, 30 min, 4 °C). Clarified lysates were incubated with TALON® metal affinity resin (Takara Bio USA) for 1 h with end-over-end rotation at 4 °C. Resin was pelleted via centrifugation (700 x *g*, 2 min, 4 °C), washed four times with equilibration buffer (400 mM NaCl, 160 mM HEPES (pH 7.5), 5 mM BME, 0.25 mM PMSF), and washed twice with wash buffer (400 mM NaCl, 160 mM HEPES (pH 7.5), 5 mM BME, 0.25 mM PMSF, 40 mM imidazole). Proteins were eluted from the resin with elution buffer (400 mM NaCl, 160 mM HEPES (pH 7.5), 5 mM BME, 0.25 mM PMSF, 400 mM imidazole). Eluted proteins were buffer-exchanged twice into equilibration buffer (400 mM NaCl, 160 mM HEPES (pH 7.5), 5 mM BME, 0.25 mM PMSF) and concentrated to ~500 μL with an Amicon® Ultra-15 Centrifugal Filter Unit (10 kDa cutoff, Millipore). Concentrated eluted proteins were cleared of protein precipitate via centrifugation (21,130 x *g*, 15 min, 4 °C). Protein concentrations were determined via Bradford assay (Bio-Rad Protein Assay Kit II; (75)), diluted with equilibration buffer (400 mM NaCl, 160 mM HEPES (pH 7.5), 5 mM BME, 0.25 mM PMSF) to the desired concentration, aliquoted into strip tubes, flash frozen in N_2_(l), and stored at −80 °C.

### Size Exclusion Chromatography

For crystallography studies, concentrated protein elutions (before the protein concentration quantification step above) were passed through a 0.22 μM filter and injected into the loading loop of a NGC Medium-Pressure Chromatography System (BioRad) fitted with a HiLoad™ 16/600 Superdex™ 75 pg column. Proteins were separated via size exclusion chromatography (SEC) using SEC buffer (50 mM NaCl, 20 mM HEPES (pH 7.5), 0.3 mM TCEP) in 1 mL fractions. Fractions from SEC were analyzed via SDS-PAGE, and the major fractions containing the target protein were pooled and then concentrated with an Amicon® Ultra-15 Centrifugal Filter Unit (10 kDa cutoff, Millipore) to ~500 mL. Concentrated SEC fractions were then dialyzed against crystal buffer (5 mM HEPES, 400 mM NaCl, 0.3 mM TCEP, pH 8.0) in a Slide-A-Lyzer™ MINI Dialysis Device (3.5 kDa cutoff, Thermo) overnight on an orbital shaker at 4 °C. Protein concentrations were determined via Bradford assay (Bio-Rad Protein Assay Kit II; Bradford 1976), diluted with crystal buffer (5 mM HEPES, 400 mM NaCl, 0.3 mM TCEP, pH 8.0) to 20 mg/mL, aliquoted into strip tubes, flash frozen in N_2_(l), and stored at −80 °C.

### Crystallization, X-ray Diffraction, and Refinement

Crystallization experiments and structure determination were conducted in the Collaborative Crystallography Core in the Department of Biochemistry at the University of Wisconsin (Madison, WI, USA). All crystallization screens and optimizations were conducted at 293K in MRC SD-2 crystallization plates, set with a STP Labtech Mosquito® crystallization robot. Hampton IndexHT and Molecular Dimensions JCSG+ were used as general screens. Crystals were cooled by direct immersion in liquid nitrogen after cryopreservation and harvest using MiTeGen MicroMounts™. X-ray experiments were conducted at the Advanced Photon Source, Argonne National Lab, GMCA@APS beamline 23ID-D. Diffraction data were collected on a Pilatus 3-6M detector and reduced using XDS (76) and XSCALE (77). Structure solution and refinement used the Phenix suite of crystallography programs (78). Iterative rounds of map fitting in Coot (79) and phenix.refine (80) were used to improve the atomic models. MOLPROBITY (81) was used the validate the structures. Data collection and refinement statistics for the structures can be found in Table S3.

The best crystals of SeMet Aim18p-Nd70-R123A grew from 26% PEG 3350, 0.2M Li_2_SO_4_, 0.1 M Na-HEPES buffer, pH 7.5. Crystals were cryopreserved by supplementing the PEG concentration to 30%. Diffraction data extending to 2.2Å was collected on 2018-02-07 at energies near the SeK-edge (peak=0.97937Å, edge=0.97961Å, and high remote at 0.96437Å). Phenix.hyss (82) located the expected 5 Se sites from one copy of the protein per asymmetric unit in space group P3_2_21. The structure was phased and traced using Phenix.autosol (83) with a phasing figure of merit=0.47, and map skew=0.6. The initial chain trace was continuous from residue 88 to the C-terminus. The final model starts at residue 83 and includes five sulfate ions per chain.

The best crystals of Aim46p-FL-WT were grown using microseeds obtained from a similar condition. The seeds were stabilized in 30% PEG 2000, 0.1M MES pH 6.5. The droplet was composed of 20 nL microseed suspension, 180 nL Aim46p-Nd62-WT, and 300 nL PEG 2000, 0.1M MES, pH 6.5. Crystals were cryoprotected with 30% PEG 2000. Data were collected on 2018-12-08 at 1.0332Å. Data extended to 2.0Å and belonged to space group P2_1_. One copy of the protein per asymmetric unit was by molecular replacement using Phaser starting from an appropriately pruned Aim18p-Nd70-R123A model (log-likelihood gain 307, translation function z-score 11.7). The final model is continuous from residue 70 to the C-terminus, and includes a ligand modeled as α-ketoglutarate.

### Chalcone Isomerase Assay

The chalcone isomerase activity assay was performed as previously described (4, 5, 7). Chalcone isomerase activity assays were run with the following reaction mixture: 150 mM NaCl, 50 mM HEPES (pH 7.5), 5% EtOH (as a cosolvent), and 200 μM naringenin chalcone (Aldlab Chemicals: AT15178). To initiate the reaction, 10 nM protein was added, mixed, and immediately run on an Infinite® M1000 multimode microplate reader (Tecan). Conversion of naringenin chalcone (yellow) into naringenin (colorless) was monitored by the depletion of absorbance at 381 nm. Absorbance measurements at 381 nm were taken every 10 s for 30 m. The resulting absorbance data were exported to Microsoft Excel and analyzed in Prism 9.2.0 (GraphPad by Dotmatics). A no protein (buffer and substrate only) control is necessary for these experiments because naringenin chalcone non-enzymatically cyclizes into naringenin at a slow rate over the course of the assay.

### Differential Scanning Fluorimetry

The general differential scanning fluorimetry (DSF) method has been documented previously (84). For screening fatty acid binding, 20 μL reactions containing 5 μM protein, 50 μM fatty acid from 10 mM stock in DMSO or equivalent vehicle (Linolenic acid [Sigma: L2376]) and Oleic acid [Sigma: O1008]), 100 mM HEPES pH 7.5, 150 mM NaCl, and 10x Sypro Orange dye (Thermo: S6651) were made in MicroAmp Optical 96-well reaction plates (Thermo: N8010560), centrifuged (200 *xg*, RT, 30 sec), and incubated at RT in the dark for 1 hour. Fluorescence was then monitored with the ROX filter using a QuantStudio 3 Real-Time PCR system along a temperature gradient from 4-95 °C at a rate of 0.025 °C/s. Protein Thermal Shift software v1.4 (Applied Biosystems) was used to determine T_m_ values by determining the maximum of first derivative curves. Melt curves flagged by the software were manually inspected. Each fatty acid-protein pair was run in triplicate, and error bars represent standard deviations. ΔT_m_ values were determined by subtracting the average of matched vehicle controls from the same plate.

### UV-Vis Spectroscopic Measurement

Purified proteins were assessed for the presence or absence of the Soret peak (Soret, 1878). 1.5 μL of 50-mg/mL purified recombinant protein were analyzed on a NanoDrop™ 2000c spectrophotometer (Thermo). Absorbance spectra from 350 nm to 700 nm (1 nm steps) were collected. The resulting absorbance spectra data were analyzed in Prism 9.2.0 (GraphPad by Dotmatics).

### Fluorometric Heme Measurement

A modified version of the fluorometry heme measurement assay was performed as described (85, 86). 100 μL of 2 M oxalic acid was added to 100-μL samples of 16 μM-purified protein or 100-μL FLAG immunoprecipitation elutions from *S. cerevisiae*. The resulting 200-μL reaction was split into two replicate 100-μL tubes (samples arrayed in a strip tube). The first replicate tube was kept in the dark at RT, and the other replicate tube was incubated at 98 °C for 30 minutes in a Mastercyler® X50s thermocycler (Eppendorph). Both replicate samples were transferred to a black-walled microplate, and fluorescence measurements (λ_ex_ = 400 nm; λ_em_ = 620 nm) were taken on an Infinite® M1000 multimode microplate reader (Tecan). The resulting fluorescence data were exported to Microsoft Excel and analyzed in Prism 9.2.0 (GraphPad by Dotmatics). Unheated samples were subtracted from the heated samples to produce the final fluorescence values and reported in relative fluorescence units (RFUs).

### Lipid Extraction from Yeast

Starter cultures (3 mL YPD) were inoculated with an individual colony of yeast and incubated (30 °C, 230 rpm, 12 h). YPGD medium (100 mL medium at ambient temperature in a sterile 250-mL Erlenmeyer flask) was inoculated with 2.5 × 10^6^ yeast cells and incubated (30 °C, 230 rpm, 18 h). Samples were harvested 18 h after inoculation, a time point that corresponds to early respiration growth. 1 × 10^8^ yeast cells were harvested by centrifugation (3,000 *x g*, 3 min, 4 °C) in screw-cap tubes, the supernatant was removed, and the cell pellet was flash frozen in N_2_(l) and stored at −80 °C. 100 μL of glass beads, and 50 μL of 150 mM KCl, and 600 μL of methanol with 0.1 μM CoQ_8_ internal standard (Avanti Polar Lipids) were added, and the cells were lysed lysed via bead beating on a Disruptor Genie® (Zymo Research) cell disruption device (2X, 5 min). To extract lipids, 400 μL of petroleum ether was added to samples and subjected to bead-beating for 3 min. Samples were then centrifuged (1000 x *g*, 2 min, 4 °C), and the ether layer (top) was transferred to a new tube. Extraction was repeated; the ether layers were combined and dried under argon gas at room temperature.

### CoQ_6_, DMQ_6_, and PPHB_6_

#### Measurement by LC-MS

Extracted dried lipids were resuspended in 50 μL of mobile phase (78% methanol, 20% isopropanol, 2% 1 M ammonium acetate pH 4.4 in water) and transferred to amber glass vials with inserts. LC-MS analysis was performed using a Thermo Vanquish Horizon UHPLC system coupled to a Thermo Exploris 240 Orbitrap mass spectrometer. For LC separation, a Vanquish binary pump system (Thermo) was used with a Waters Acquity CSH C18 column (100 mm × 2.1 mm, 1.7 mm particle size) held at 35 °C under 300 μL/min flow rate. Mobile phase A consisted of 5 mM ammonium acetate in ACN:H_2_O (70:30, v/v) containing 125 μL/L acetic acid. Mobile phase B consisted of 5 mM ammonium acetate in IPA:ACN (90:10, v/v) with the same additive. For each sample run, mobile phase B was initially held at 2% for 2 min and then increased to 30% over 3 min. Mobile phase B was further increased to 50% over 1 min and 85% over 14 min and then raised to 99% over 1 min and held for 4 min. The column was re-equilibrated for 5 min at 2% B before the next injection. 5 μL of sample were injected by a Vanquish Split Sampler HT autosampler (Thermo Scientific), while the autosampler temperature was kept at 4 °C. The samples were ionized by a heated ESI source kept at a vaporizer temperature of 350 °C. Sheath gas was set to 50 units, auxiliary gas to 8 units, sweep gas to 1 unit, and the spray voltage was set to 3,500 V for positive mode and 2,500 V for negative mode. The inlet ion transfer tube temperature was kept at 325 °C with 70% RF lens. For targeted analysis, the MS was operated in positive parallel reaction monitoring (PRM) mode with polarity switching acquiring scheduled, targeted scans to CoQ intermediates: PPHB, DMQ, and CoQ. MS acquisition parameters include resolution of 15,000, stepped HCD collision energy (25%, 30% for positive mode and 20%, 40%, 60% for negative mode), and 3 s dynamic exclusion. Automatic gain control (AGC) targets were set to standard mode.

#### Data Analysis

The resulting CoQ intermediate (oxidized form) data were processed using TraceFinder 5.1 (Thermo).

#### Dali Server Protein Structure Comparison

The crystal structures of Aim18p-Nd70-R123A and Aim46p-Nd62-WT were submitted as query protein structures to the Dali server (http://ekhidna2.biocenter.helsinki.fi/dali/; (28)) and compared against all of the structures in the Protein Data Bank (PDB). PDB protein structural matches were provided in a list sorted by Z-score.

#### PyMOL™ Protein Structure Visualization

Protein crystal structures (pdb format) were imported into the PyMOL™ 2.4.0 software. Proteins were structurally aligned using the super command and colored for visual clarity. In some cases, regions of proteins were made slightly transparent to highlight areas of importance. Scenes were exported using the ray command (specifically, ray_trace_mode,3) as png files and annotated in Adobe Illustrator 26.5.

#### Statistical analysis

Statistical analyses were calculated with Microsoft Excel and visualized with Prism 9.2.0 (GraphPad by Dotmatics). For all experiments, “mean” indicates the arithmetic mean, SD indicates the standard deviation, and “*n*” indicates independent biological or technical replicates, as specified. *P*-values were calculated using a two-tailed, unpaired Student’s t-test (for significance reporting: n.s. = *p* > 0.05; * = *p* ≤ 0.05; ** = p ≤ 0.01; *** = p ≤ 0.001).

## Supporting information

Tabl2 S2

## DATA AVAILABILITY

All data are contained within the article or supporting information. The structures presented in this article have been deposited in the Protein Data Bank (PDB) with the codes 8EW8 (Aim18p-Nd70-R123A) and 8EW9 (Aim46p-Nd62-WT).

## SUPPORTING INFORMATION

This article contains supporting information (7, 15, 16, 69–73).

## ACKNOWLEDGEMENTS

We thank Jared Rutter for providing the parental W303-1A yeast strain used in this study, Nora Vögtle and Betty Craig for mitochondrial antibodies, Betty Craig for use of their ultracentrifuge, Linda Horianopoulos for assistance with western blot densitometry, the Hittinger and Donohue labs and the for the use of their lab spaces, and the former and current members of the Pagliarini and Hittinger labs for their scientific insight and feedback during the course of this study.

## FUNDING AND ADDITIONAL INFORMATION

This work was financially supported by the NIH awards R35 GM131795 and P41 GM108538 and funds from the BJC Investigator Program (to D.J.P.). Research in the Hittinger Lab is supported by the National Science Foundation under Grant Nos. DEB-1442148 and DEB-2110403, in part by the DOE Great Lakes Bioenergy Research Center (DOE BER Office of Science DE-SC0018409), the United States Department of Agriculture National Institute of Food and Agriculture (Hatch Project 1020204), and the Office of the Vice Chancellor for Research and Graduate Education with funding from the Wisconsin Alumni Research Foundation (H. I. Romnes Faculty Fellow). John Wolters received support from National Institutes of Health (T32 HG002760-16) and National Science Foundation (Postdoctoral Research Fellowship in Biology 1907278). Nathan Murray received support from the Chemistry-Biology Interface Training Program (NIH T32GM008505) and NSF GRFP (DGE-1747503). Crystallization, data collection, and structure solution and refinement were conducted by the Department of Biochemistry Collaborative Crystallography Core at the University of Wisconsin, which is supported by user fees and the department. This research used resources of the Advanced Photon Source, a U.S. Department of Energy (DOE) Office of Science User Facility operated for the DOE Office of Science by Argonne National Laboratory under Contract No. DE-AC02-06CH11357. GM/CA@APS has been funded by the National Cancer Institute (ACB-12002) and the National Institute of General Medical Sciences (AGM-12006, P30GM138396). The content is solely the responsibility of the authors and does not necessarily represent the official views of the National Institutes of Health, National Science Foundation, Department of Energy, National Cancer Institute, National Institute of General Medical Sciences, Department of Agriculture, or Wisconsin Alumni Research Foundation.

## ABBREVIATIONS AND NOMENCLATURE

The abbreviations used in this study are:

CHI: chalcone isomerase
CHIL: chalone isomerase like
FAP: fatty acid binding protein
DNA: deoxyribonucleic acid
HMG-CoA: β-Hydroxy β-methylglutaryl-CoA
Ndxx: N-terminal deletion of xx amino acids
ORF: open reading frame
SC: synthetic complete
Ura-: uracil drop-out media
GPD: glyceraldehyde-3-phosphate dehydrogenase
SS: salmon sperm DNA
PEG: polyethylene glycol
FLAG: immunoprecipitation epitope consisting of Asp-Tyr-Lys-Asp-Asp-Asp-Asp-Lys
bp: base pairs
SDM: site-directed mutagenesis
BME: β-mercaptoethanol
TCA: trichloroacetic acid
LDS: lithium dodecyl sulfate
SDS-PAGE: sodium dodecyl sulfate–polyacrylamide gel electrophoresis
PVDF: polyvinylidene fluoride
NFDM: non-fat dry milk
TBST: tris buffered saline plus 0.05% tween-20 (v/v)
(v/v): percent volume per 100 mL volume
(w/v): percent weight per 100 mL volume
GAPDH: glyceraldehyde-3-phosphate dehydrogenase
CYTB: cytochrome b
CIT: citrate synthase
TOM: translocase of the outer membrane
TIM: translocase of the inner membrane
SEC: size-exclusion chromatography
TCEP: tris(2-carboxyethyl)phosphine
DTT: dithiothreitol
DSF: differential scanning fluorimetry
UV-Vis: ultraviolet–visible spectroscopy
CoQ: coenzyme Q or ubiquinone
DMQ: demethoxyubiquinone
PPHB: polyprenyl-4-hydroxybenzoate
PDB: Protein Data Bank

## SUPPLEMENTAL FIGURES

**Figure S1.**
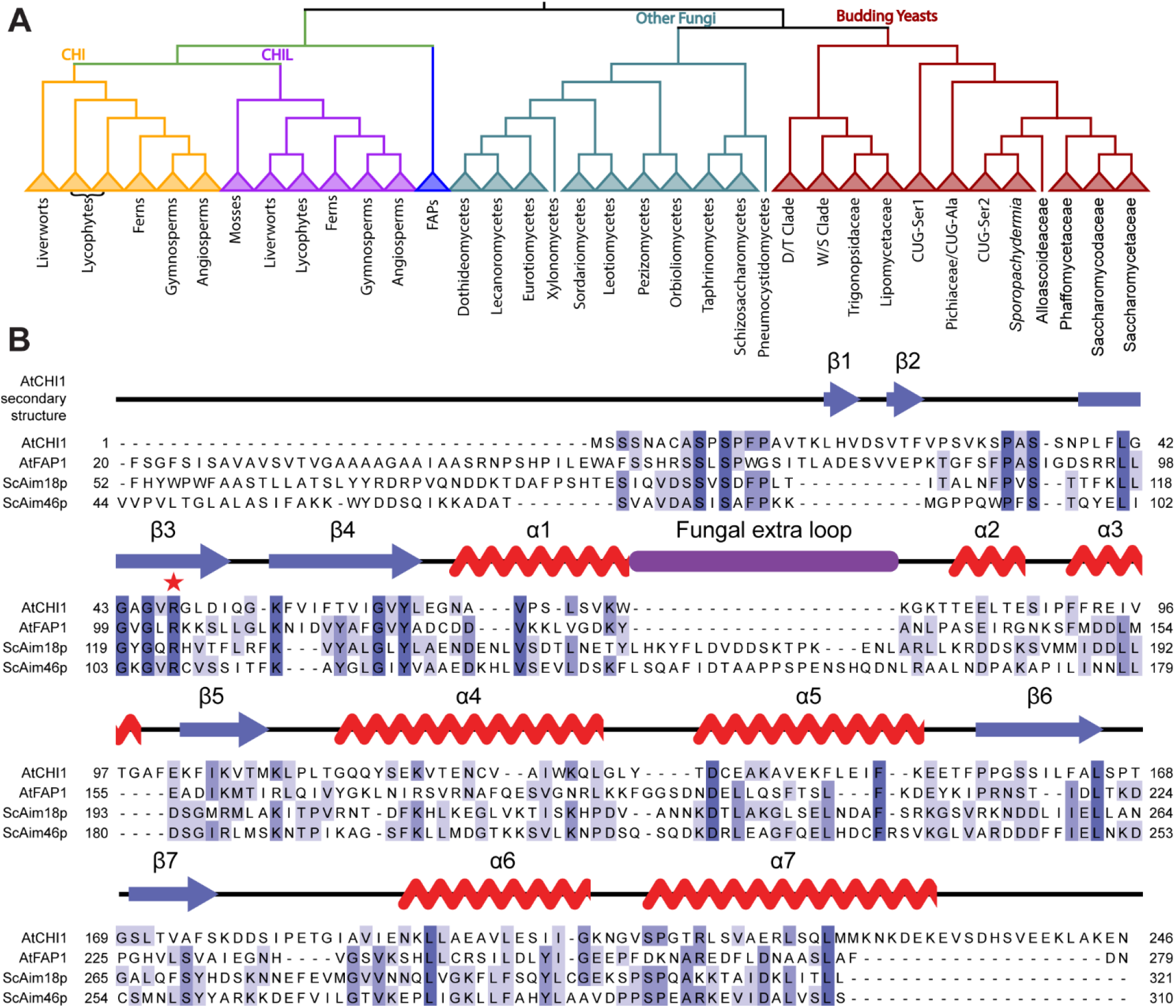
Aim18p and Aim46p are yeast proteins with sequence homology to plant CHIs. **A**. Phylogenetic tree of Fig. 1B, built with all species, but collapsed to a visualizable number of clades. Full Newick tree file and additional information for all sequences included for the phylogenetic tree presented in Fig. S1A are included in Supporting Documents S1.zip. Aside from gene duplications, the CHI topology generally followed the species phylogeny, although some lineages were incorrectly split, often by poorly supported branches. Plant sequencies diverged early into multiple groups with many species possessing multiple CHI and CHIL sequences, whereas nearly all budding yeast species contain only a single CHI homolog. **B**. Protein alignment of AtCHI1, AtFAP1, Aim18p, and Aim46p protein sequences. Input protein sequences for Aim18p (P38884), Aim46p (P38885), AtCHI1 (P41088), and AtFAP1 (Q9M1X2) were obtained from (https://uniprot.org/). AtFAP1, Aim18p, and Aim46p sequences were truncated to remove organelle targeting sequences. Secondary protein structure of AtCHI1 crystal structure (PDB: 4DOI) above corresponding regions of the alignment. Sequences colored according to %identity in shades of blue. Conserved CHI catalytic arginine indicated with a red star. Sequence and alignment information for the alignment presented in Fig. S1B are included in Supporting Documents S1.zip.

**Figure S2.**
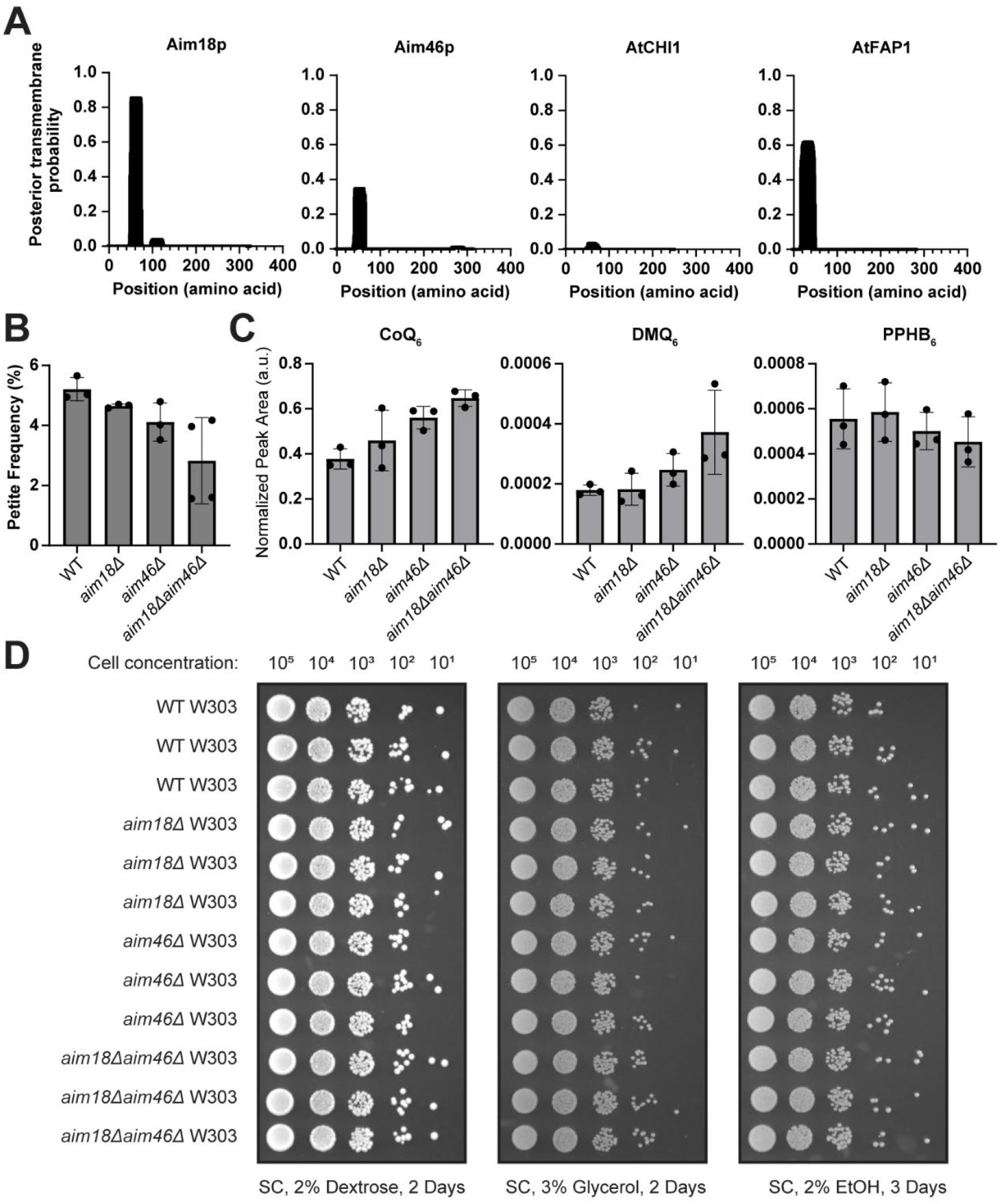
Aim18p and Aim46p are inner mitochondrial membrane proteins. **A**. Phobius plots indicating the presence of an N-terminal transmembrane domain for Aim18p, Aim46p, and AtFAP1. Input protein sequences for Aim18p (P38884), Aim46p (P38885), AtCHI1 (P41088), and AtFAP1 (Q9M1X2) were obtained from (https://uniprot.org/). Phobius data retrieved from (https://phobius.sbc.su.se/) and visualized in Prism 9.2.0 (GraphPad by Dotmatics). **B**. Yeast petite frequency assay of WT, *aim18Δ, aim46Δ*, and *aim18Δaim46Δ* W303-1A grown for 2-3 days on YPDG plates containing 0.1% glucose (w/v) and 3% glycerol (w/v). *n* = 3 independent biological replicates for WT, *aim18Δ*, and *aim46Δ* W303-1A. *n* = 4 independent biological replicates for *aim18Δaim46Δ* W303-1A. **C**. CoQ and CoQ intermediates extracted from WT, *aim18Δ, aim46Δ*, and *aim18Δaim46Δ* W303-1A yeast and measured by LC-MS. *n* = 3 independent biological replicates. **D**. Yeast drop assay of WT, *aim18Δ, aim46Δ*, and *aim18Δaim46Δ* W303-1A grown for 2-3 days on SC media containing 2% glucose (w/v), 3% glycerol (w/v), and 2% EtOH (v/v). *n* = 3 independent biological replicates.

**Figure S3.**
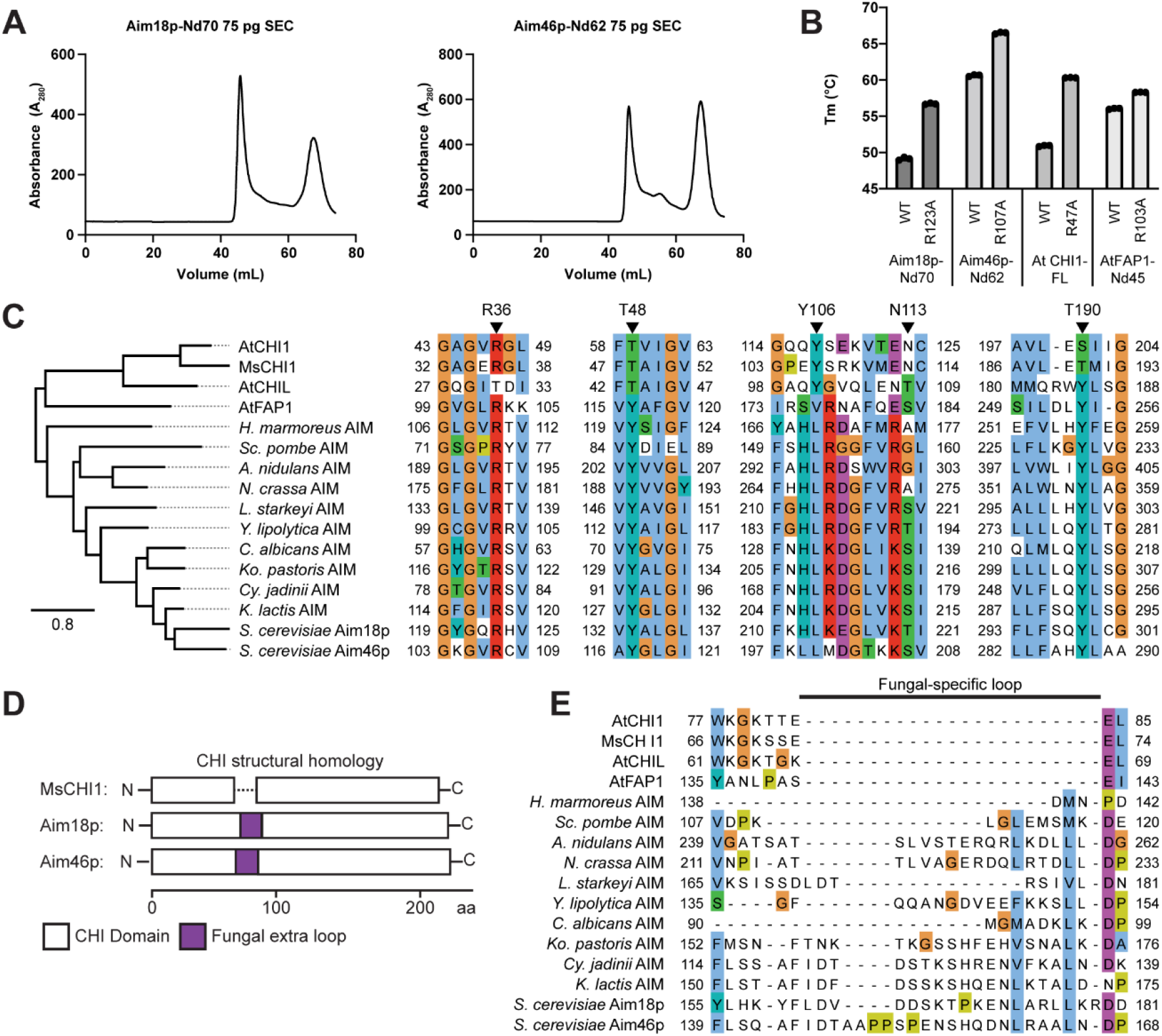
Aim18p and Aim46p adopt the CHI-fold but lack CHI-fold activities. **A**. Size exclusion chromatography peaks from purified recombinant Aim18-Nd70-WT and Aim46-Nd62-WT. Recombinant proteins were separated by size on an NGC Medium-Pressure Chromatography System (BioRad) fitted with a HiLoad™ 16/600 Superdex™ 75 pg column, and 1-mL fractions were collected. The void volume for this column was ~44 mL, while 30 kDa proteins elute at a volume of ~70 mL. A_280_ measurements were continuously taken to monitor relative protein concentration. **B.** Thermal melting temperatures (T_m_) of CHI-family proteins (mean ± SD, *n* = 3 independent technical replicates). **C**. Phylogenetic tree and protein sequence alignment of AtCHI1, MsCHI1, AtCHIL, AtFAP1, and fungal CHI-domain containing proteins. Input protein sequences for Aim18p (P38884), Aim46p (P38885), AtCHI1 (P41088), and AtFAP1 (Q9M1X2) were obtained from (https://uniprot.org/). Input protein sequences for fungal CHI-domain containing proteins were obtained from the translated gBlocks in Table S2. The five key CHI active site residues of MsCHI1 are indicated with arrows and refer to the protein sequence of MsCHI1 from *Medicago sativa*. Note that the AIM protein phylogeny matches the fungal species phylogeny (16) with the exception of *Sc. pombe*, which is an outgroup to the other ascomycetes; *H. marmoreus* is a basidiomycete. Phylogenetic tree branch lengths are proportional to the number of substitutions per site. Sequences colored using the Clustal color palette by conservation. **D**. Diagram of structural homology of Aim18p and Aim46p compared to MsCHI1. White boxes indicate regions of high CHI domain structural homology. Purple boxes indicate the extra loop only present in Aim18p and Aim46p. **E.** Subsection of the protein sequence alignment presented in Fig. S3C highlighting the fungal-specific loop comparison between fungal and plant CHI-domain-containing proteins. Sequences colored using the Clustal color palette by conservation. Full Newick tree file and additional information for all sequences included for the phylogenetic tree presented in Fig. S3 are included in Supporting Documents S1.zip.

**Figure S4.**
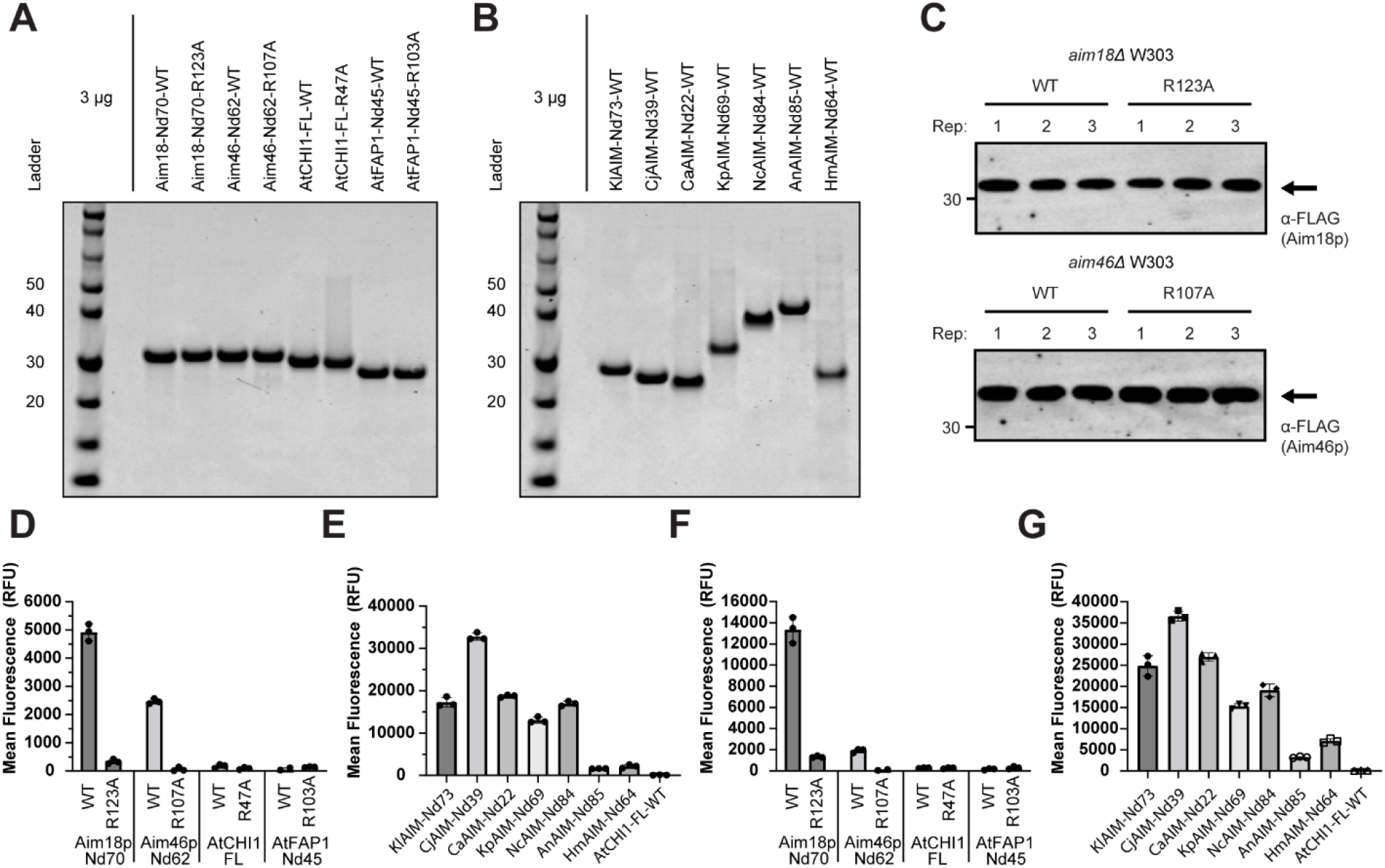
Aim18p and Aim46p are hemoproteins. **A**. SDS-PAGE analysis of 3-μg elutions of purified recombinant Aim18p-Nd70, Aim46p-Nd62, AtCHI1, and AtFAP1-Nd45 variants. **B**. SDS-PAGE analysis of purified recombinant fungal AIM homologs. Recombinant fungal AIM homologs are named based on their species of origin (KlAIM = *K. lactis*; CjAIM = *Cy. jadinii*; CaAIM = *C. albicans*; KpAIM = *Ko. pastoris*; NcAIM = *N. crassa*; AnAIM = *A. nidulans*; HmAIM = *H. marmoreus*). **C**. Western blot analysis of 0.5% immunoprecipitation elution fractions from heme measurements in Fig. 4C. **D**. Fluorometric oxalic acid heme determination assay with 16-μM recombinant Aim18p-Nd70, Aim46-Nd62, AtCHI1, and AtFAP1 proteins purified in Fig. 4A (mean ± SD, *n* = 3 independent technical replicates). **E**. Fluorometric oxalic acid heme determination assay with 16-μM recombinant fungal AIM homolog proteins purified in Fig. 4A (mean ± SD, *n* = 3 independent technical replicates). **F**. Fluorometric oxalic acid heme determination assay with 16-μM recombinant Aim18p-Nd70, Aim46-Nd62, AtCHI1, and AtFAP1 proteins purified in Fig. 4A (mean ± SD, *n* = 3 independent technical replicates). Data are representative of a second independent protein purification of proteins in Fig. S4D. **G**. Fluorometric oxalic acid heme determination assay with 16-μM recombinant fungal AIM homolog proteins purified in Fig. 4A (mean ± SD, *n* = 3 independent technical replicates). Data are representative of a second independent protein purification of proteins in Fig. S4E.

## REFERENCES

1. Shimokoriyama, M. (1957) Interconversion of Chalcones and Flavanones of a Phloroglucinol-type Structure. J Am Chem Soc. 79, 4199–4202

2. Jez, J. M., Bowman, M. E., Dixon, R. A., and Noel, J. P. (2000) Structure and mechanism of the evolutionarily unique plant enzyme chalcone isomerase. Nature Structural Biology 2000 7:9. 7, 786–791

3. Nishihara, M., Nakatsuka, T., and Yamamura, S. (2005) Flavonoid components and flower color change in transgenic tobacco plants by suppression of chalcone isomerase gene. FEBS Lett. 579, 6074–6078

4. Jez, J. M., and Noel, J. P. (2002) Reaction mechanism of chalcone isomerase: pH dependence, diffusion control, and product binding differences. Journal of Biological Chemistry. 277, 1361–1369

5. Jez, J. M., Bowman, M. E., and Noel, J. P. (2002) Role of hydrogen bonds in the reaction mechanism of chalcone isomerase. Biochemistry. 41, 5168–5176

6. Ngaki, M. N., Louie, G. v., Philippe, R. N., Manning, G., Pojer, F., Bowman, M. E., Li, L., Larsen, E., Wurtele, E. S., and Noel, J. P. (2012) Evolution of the chalcone-isomerase fold from fatty-acid binding to stereospecific catalysis. Nature. 485, 530–533

7. Kaltenbach, M., Burke, J. R., Dindo, M., Pabis, A., Munsberg, F. S., Rabin, A., Kamerlin, S. C. L., Noel, J. P., and Tawfik, D. S. (2018) Evolution of chalcone isomerase from a noncatalytic ancestor article. Nat Chem Biol. 14, 548–555

8. Gensheimer, M. (2004) Chalcone isomerase family and fold: No longer unique to plants. Protein Science. 13, 540–544

9. Dastmalchi, M., and Dhaubhadel, S. (2015) Soybean chalcone isomerase: evolution of the fold, and the differential expression and localization of the gene family. Planta. 241, 507–523

10. Sobreira, A. C. M., Pinto, F. das C. L., Florêncio, K. G. D., Wilke, D. v., Staats, C. C., Streit, R. de A. S., Freire, F. das C. de O., Pessoa, O. D. L., Trindade-Silva, A. E., and Canuto, K. M. (2018) Endophytic fungus Pseudofusicoccum stromaticum produces cyclopeptides and plant-related bioactive rotenoids. RSC Adv. 8, 35575–35586

11. Cheng, A. X., Zhang, X., Han, X. J., Zhang, Y. Y., Gao, S., Liu, C. J., and Lou, H. X. (2018) Identification of chalcone isomerase in the basal land plants reveals an ancient evolution of enzymatic cyclization activity for synthesis of flavonoids. New Phytologist. 217, 909–924

12. Yin, Y. chao, Zhang, X. dong, Gao, Z. qiang, Hu, T., and Liu, Y. (2019) The Research Progress of Chalcone Isomerase (CHI) in Plants. Mol Biotechnol. 61, 32–52

13. Wan, Q., Bai, T., Liu, M., Liu, Y., Xie, Y., Zhang, T., Huang, M., and Zhang, J. (2022) Comparative Analysis of the Chalcone-Flavanone Isomerase Genes in Six Citrus Species and Their Expression Analysis in Sweet Orange (Citrus sinensis). Front Genet. 13, 734

14. Saito, K., Yonekura-Sakakibara, K., Nakabayashi, R., Higashi, Y., Yamazaki, M., Tohge, T., and Fernie, A. R. (2013) The flavonoid biosynthetic pathway in Arabidopsis: Structural and genetic diversity. Plant Physiology and Biochemistry. 72, 21–34

15. Shen, X. X., Opulente, D. A., Kominek, J., Zhou, X., Steenwyk, J. L., Buh, K. v., Haase, M. A. B., Wisecaver, J. H., Wang, M., Doering, D. T., Boudouris, J. T., Schneider, R. M., Langdon, Q. K., Ohkuma, M., Endoh, R., Takashima, M., Manabe, R. ichiroh, Čadež, N., Libkind, D., Rosa, C. A., DeVirgilio, J., Hulfachor, A. B., Groenewald, M., Kurtzman, C. P., Hittinger, C. T., and Rokas, A. (2018) Tempo and Mode of Genome Evolution in the Budding Yeast Subphylum. Cell. 175, 1533–1545.e20

16. Li, Y., Steenwyk, J. L., Chang, Y., Wang, Y., James, T. Y., Stajich, J. E., Spatafora, J. W., Groenewald, M., Dunn, C. W., Hittinger, C. T., Shen, X. X., and Rokas, A. (2021) A genome-scale phylogeny of the kingdom Fungi. Curr Biol. 31, 1653–1665.e5

17. Fukasawa, Y., Tsuji, J., Fu, S. C., Tomii, K., Horton, P., and Imai, K. (2015) MitoFates: improved prediction of mitochondrial targeting sequences and their cleavage sites. Mol Cell Proteomics. 14, 1113–1126

18. Käll, L., Krogh, A., and Sonnhammer, E. L. L. (2007) Advantages of combined transmembrane topology and signal peptide prediction-the Phobius web server. Nucleic Acids Res. 10.1093/nar/gkm256

19. Huh, W. K., Falvo, J. v., Gerke, L. C., Carroll, A. S., Howson, R. W., Weissman, J. S., and O’Shea, E. K. (2003) Global analysis of protein localization in budding yeast. Nature 2003 425:6959. 425, 686–691

20. Reinders, J., Zahedi, R. P., Pfanner, N., Meisinger, C., and Sickmann, A. (2006) Toward the complete yeast mitochondrial proteome: Multidimensional separation techniques for mitochondrial proteomics. J Proteome Res. 5, 1543–1554

21. Vögtle, F. N., Burkhart, J. M., Gonczarowska-Jorge, H., Kücükköse, C., Taskin, A. A., Kopczynski, D., Ahrends, R., Mossmann, D., Sickmann, A., Zahedi, R. P., and Meisinger, C. (2017) Landscape of submitochondrial protein distribution. Nat Commun. 10.1038/s41467-017-00359-0

22. Morgenstern, M., Stiller, S. B., Lübbert, P., Peikert, C. D., Dannenmaier, S., Drepper, F., Weill, U., Höß, P., Feuerstein, R., Gebert, M., Bohnert, M., van der Laan, M., Schuldiner, M., Schütze, C., Oeljeklaus, S., Pfanner, N., Wiedemann, N., and Warscheid, B. (2017) Definition of a High-Confidence Mitochondrial Proteome at Quantitative Scale. Cell Rep. 19, 2836–2852

23. Zahedi, R. P., Sickmann, A., Boehm, A. M., Winkler, C., Zufall, N., Schö, B., Guiard, B., Pfanner, N., and Meisinger, C. (2006) Proteomic Analysis of the Yeast Mitochondrial Outer Membrane Reveals Accumulation of a Subclass of Preproteins □ D. Mol Biol Cell. 17, 1436–1450

24. Kim, H., Botelho, S. C., Park, K., and Kim, H. (2015) Use of carbonate extraction in analyzing moderately hydrophobic transmembrane proteins in the mitochondrial inner membrane. Protein Sci. 24, 2063

25. Hess, D. C., Myers, C., Huttenhower, C., Hibbs, M. A., Hayes, A. P., Paw, J., Clore, J. J., Mendoza, R. M., Luis, B. S., Nislow, C., Giaever, G., Costanzo, M., Troyanskaya, O. G., and Caudy, A. A. (2009) Computationally driven, quantitative experiments discover genes required for mitochondrial biogenesis. PLoS Genet. 10.1371/journal.pgen.1000407

26. Stefely, J. A., Kwiecien, N. W., Freiberger, E. C., Richards, A. L., Jochem, A., Rush, M. J. P., Ulbrich, A., Robinson, K. P., Hutchins, P. D., Veling, M. T., Guo, X., Kemmerer, Z. A., Connors, K. J., Trujillo, E. A., Sokol, J., Marx, H., Westphall, M. S., Hebert, A. S., Pagliarini, D. J., and Coon, J. J. (2016) Mitochondrial protein functions elucidated by multi-omic mass spectrometry profiling. Nat Biotechnol. 34, 1191–1197

27. Jacques-Louis Soret (1883) Analyse spectrale: Sur le spectre d’absorption du sang dans la partie violette et ultra-violette. Comptes rendus de l’Académie des sciences (In French). 97, 1269–1270

28. Holm, L. (2022) Dali server: structural unification of protein families. Nucleic Acids Res. 50, W210–W215

29. Abendroth, J., Buchko, G. W., Liew, F. N., Nguyen, J. N., Kim, H. J., and Kim, H. J. (2022) Structural Characterization of Cytochrome c’β-Met from an Ammonia-Oxidizing Bacterium. Biochemistry. 61, 563–574

30. Ralston, L., Subramanian, S., Matsuno, M., and Yu, O. (2005) Partial Reconstruction of Flavonoid and Isoflavonoid Biosynthesis in Yeast Using Soybean Type I and Type II Chalcone Isomerases 1[w]. Plant Physiol. 137, 1375–1388

31. Morita, Y., Takagi, K., Fukuchi-Mizutani, M., Ishiguro, K., Tanaka, Y., Nitasaka, E., Nakayama, M., Saito, N., Kagami, T., Hoshino, A., and Iida, S. (2014) A chalcone isomerase-like protein enhances flavonoid production and flower pigmentation. Plant J. 78, 294–304

32. Xu, H., Lan, Y., Xing, J., Li, Y., Liu, L., and Wang, Y. (2022) AfCHIL, a Type IV Chalcone Isomerase, Enhances the Biosynthesis of Naringenin in Metabolic Engineering. Front Plant Sci. 13, 1458

33. Paoli, M., Marles-wright, J., and Smith, A. (2002) Structure-function relationships in heme-proteins. DNA Cell Biol. 21, 271–280

34. Poulos, T. L. (2007) The Janus nature of heme. Nat Prod Rep. 24, 504–510

35. Orengo, C. A., and Thornton, J. M. (1993) Alpha plus beta folds revisited: some favoured motifs. Structure. 1, 105–120

36. Reedy, C. J., and Gibney, B. R. (2004) Heme Protein Assemblies. Chem Rev. 104, 617–649

37. Li, T., Bonkovsky, H. L., and Guo, J. T. (2011) Structural analysis of heme proteins: Implications for design and prediction. BMC Struct Biol. 10.1186/1472-6807-11-13

38. Taylor, B. L., and Zhulin, I. B. (1999) PAS Domains: Internal Sensors of Oxygen, Redox Potential, and Light. Microbiology and Molecular Biology Reviews. 63, 479

39. Gilles-Gonzalez, M. A., and Gonzalez, G. (2004) Signal transduction by heme-containing PAS-domain proteins. J Appl Physiol. 96, 774–783

40. Delgado-Nixon, V. M., Gonzalez, G., and Gilles-Gonzalez, M. A. (2000) Dos, a heme-binding PAS protein from Escherichia coli, is a direct oxygen sensor. Biochemistry. 39, 2685–2691

41. Yamawaki, T., Mizuno, M., Ishikawa, H., Takemura, K., Kitao, A., Shiro, Y., and Mizutani, Y. (2021) Regulatory Switching by Concerted Motions on the Microsecond Time Scale of the Oxygen Sensor Protein FixL. Journal of Physical Chemistry B. 125, 6847–6856

42. Izadi, N., Henry, Y., Haladjian, J., Goldberg, M. E., Wandersman, C., Delepierre, M., and Lecroisey, A. (1997) Purification and characterization of an extracellular heme-binding protein, HasA, involved in heme iron acquisition. Biochemistry. 36, 7050–7057

43. Arnoux, P., Haser, R., Izadi, N., Lecroisey, A., Delepierre, M., Wandersman, C., and Czjzek, M. (1999) The crystal structure of HasA, a hemophore secreted by Serratia marcescens. Nature Structural Biology 1999 6:6. 6, 516–520

44. Goodfellow, B. J., Freire, F., Carvalho, A. L., Aveiro, S. S., Charbonnier, P., Moulis, J. M., Delgado, L., Ferreira, G. C., Rodrigues, J. E., Poussin-Courmontagne, P., Birck, C., McEwen, A., and Macedo, A. L. (2021) The SOUL family of heme-binding proteins: Structure and function 15 years later. Coord Chem Rev. 448, 214189

45. Acharya, G., Kaur, G., and Subramanian, S. (2016) Evolutionary relationships between heme-binding ferredoxin α+β barrels. BMC Bioinformatics. 17, 1–11

46. Hofbauer, S., Pfanzagl, V., Michlits, H., Schmidt, D., Obinger, C., and Furtmüller, P. G. (2021) Understanding molecular enzymology of porphyrin-binding α+β barrel proteins - One fold, multiple functions. Biochimica et Biophysica Acta (BBA) - Proteins and Proteomics. 1869, 140536

47. Edwards, A. M., Isserlin, R., Bader, G. D., Frye, S. v., Willson, T. M., and Yu, F. H. (2011) Too many roads not taken. Nature 2011 470:7333. 470, 163–165

48. Kustatscher, G., Collins, T., Gingras, A. C., Guo, T., Hermjakob, H., Ideker, T., Lilley, K. S., Lundberg, E., Marcotte, E. M., Ralser, M., and Rappsilber, J. (2022) Understudied proteins: opportunities and challenges for functional proteomics. Nature Methods 2022 19:7. 19, 774–779

49. Yang, J., and Zhang, Y. (2015) I-TASSER server: new development for protein structure and function predictions. Web Server issue Published online. 10.1093/nar/gkv342

50. Zheng, W., Zhang, C., Li, Y., Pearce, R., Bell, E. W., and Zhang, Y. (2021) Folding non-homologous proteins by coupling deep-learning contact maps with I-TASSER assembly simulations. Cell Reports Methods. 1, 100014

51. Jumper, J., Evans, R., Pritzel, A., Green, T., Figurnov, M., Ronneberger, O., Tunyasuvunakool, K., Bates, R., Žídek, A., Potapenko, A., Bridgland, A., Meyer, C., Kohl, S. A. A., Ballard, A. J., Cowie, A., Romera-Paredes, B., Nikolov, S., Jain, R., Adler, J., Back, T., Petersen, S., Reiman, D., Clancy, E., Zielinski, M., Steinegger, M., Pacholska, M., Berghammer, T., Bodenstein, S., Silver, D., Vinyals, O., Senior, A. W., Kavukcuoglu, K., Kohli, P., and Hassabis, D. (2021) Highly accurate protein structure prediction with AlphaFold. Nature 2021 596:7873. 596, 583–589

52. Keskin, O., and Nussinov, R. (2005) Favorable scaffolds: proteins with different sequence, structure and function may associate in similar ways. Protein Engineering, Design and Selection. 18, 11–24

53. Medvedevid, K. E., Kinch, L. N., Schaeffer, R. D., and Grishin, N. v. (2019) Functional analysis of Rossmann-like domains reveals convergent evolution of topology and reaction pathways. PLoS Comput Biol. 10.1371/JOURNAL.PCBI.1007569

54. Adams, H. R., Krewson, C., Vardanega, J. E., Fujii, S., Moreno, T., Chicano, Sambongi, Y., Svistunenko, D., Paps, J., Andrew, C. R., and Hough, M. A. (2019) One fold, two functions: cytochrome P460 and cytochrome c ‘-β from the methanotroph Methylococcus capsulatus (Bath). Chem Sci. 10, 3031–3041

55. Katoh, K., and Toh, H. (2010) Parallelization of the MAFFT multiple sequence alignment program. Bioinformatics. 26, 1899–1900

56. Capella-Gutiérrez, S., Silla-Martínez, J. M., and Gabaldón, T. (2009) trimAl: a tool for automated alignment trimming in large-scale phylogenetic analyses. Bioinformatics. 25, 1972–1973

57. Price, M. N., Dehal, P. S., and Arkin, A. P. (2010) FastTree 2 – Approximately Maximum-Likelihood Trees for Large Alignments. PLoS One. 5, e9490

58. Letunic, I., and Bork, P. (2021) Interactive Tree Of Life (iTOL) v5: an online tool for phylogenetic tree display and annotation. Nucleic Acids Res. 49, W293–W296

59. Martin, D. P., Murrell, B., Golden, M., Khoosal, A., and Muhire, B. (2015) RDP4: Detection and analysis of recombination patterns in virus genomes. Virus Evol. 10.1093/VE/VEV003

60. Waterhouse, A. M., Procter, J. B., Martin, D. M. A., Clamp, M., and Barton, G. J. (2009) Jalview Version 2--a multiple sequence alignment editor and analysis workbench. Bioinformatics. 25, 1189–1191

61. Dereeper, A., Guignon, V., Blanc, G., Audic, S., Buffet, S., Chevenet, F., Dufayard, J. F., Guindon, S., Lefort, V., Lescot, M., Claverie, J. M., and Gascuel, O. (2008) Phylogeny.fr: robust phylogenetic analysis for the non-specialist. Nucleic Acids Res. 36, W469

62. Edgar, R. C. (2004) MUSCLE: multiple sequence alignment with high accuracy and high throughput. Nucleic Acids Res. 32, 1792

63. Guindon, S., Dufayard, J. F., Lefort, V., Anisimova, M., Hordijk, W., and Gascuel, O. (2010) New Algorithms and Methods to Estimate Maximum-Likelihood Phylogenies: Assessing the Performance of PhyML 3.0. Syst Biol. 59, 307–321

64. Anisimova, M., and Gascuel, O. (2006) Approximate likelihood-ratio test for branches: A fast, accurate, and powerful alternative. Syst Biol. 55, 539–552

65. Chevenet, F., Brun, C., Bañuls, A. L., Jacq, B., and Christen, R. (2006) TreeDyn: Towards dynamic graphics and annotations for analyses of trees. BMC Bioinformatics. 7, 1–9

66. Baudin, A., Ozier-kalogeropoulos, O., Denouel, A., Lacroute, F., and Cullin, C. (1993) A simple and efficient method for direct gene deletion in Saccharomyces cerevisiae. Nucleic Acids Res. 21, 3329–3330

67. Meisinger, C., Pfanner, N., and Truscott, K. N. (2006) Isolation of yeast mitochondria. Methods Mol Biol. 313, 33–39

68. Schneider, C. A., Rasband, W. S., and Eliceiri, K. W. (2012) NIH Image to ImageJ: 25 years of image analysis. Nature Methods 2012 9:7. 9, 671–675

69. Vögtle, F. N., Brändl, B., Larson, A., Pendziwiat, M., Friederich, M. W., White, S. M., Basinger, A., Kücükköse, C., Muhle, H., Jähn, J. A., Keminer, O., Helbig, K. L., Delto, C. F., Myketin, L., Mossmann, D., Burger, N., Miyake, N., Burnett, A., van Baalen, A., Lovell, M. A., Matsumoto, N., Walsh, M., Yu, H. C., Shinde, D. N., Stephani, U., van Hove, J. L. K., Müller, F. J., and Helbig, I. (2018) Mutations in PMPCB Encoding the Catalytic Subunit of the Mitochondrial Presequence Protease Cause Neurodegeneration in Early Childhood. The American Journal of Human Genetics. 102, 557–573

70. Voisine, C., Cheng, Y. C., Ohlson, M., Schilke, B., Hoff, K., Beinert, H., Marszalek, J., and Craig, E. A. (2001) Jac1, a mitochondrial J-type chaperone, is involved in the biogenesis of Fe/S clusters in Saccharomyces cerevisiae. Proceedings of the National Academy of Sciences. 98, 1483–1488

71. D’Silva, P. R., Schilke, B., Hayashi, M., and Craig, E. A. (2008) Interaction of the J-protein heterodimer Pam18/Pam16 of the mitochondrial import motor with the translocon of the inner membrane. Mol Biol Cell. 19, 424–432

72. Liu, Q., Krzewska, J., Liberek, K., and Craig, E. A. (2001) Mitochondrial Hsp70 Ssc1: Role in Protein Folding. Journal of Biological Chemistry. 276, 6112–6118

73. Guo, X., Niemi, N. M., Hutchins, P. D., Condon, S. G. F., Jochem, A., Ulbrich, A., Higbee, A. J., Russell, J. D., Senes, A., Coon, J. J., and Pagliarini, D. J. (2017) Ptc7p Dephosphorylates Select Mitochondrial Proteins to Enhance Metabolic Function. Cell Rep. 18, 307–313

74. Fox, B. G., and Blommel, P. G. (2009) Autoinduction of protein expression. Curr Protoc Protein Sci. 10.1002/0471140864.PS0523S56

75. Bradford, M. (1976) A rapid and sensitive method for the quantitation of microgram quantities of protein utilizing the principle of protein-dye binding. Anal Biochem. 72, 248–254

76. Kabsch, W. (2010) XDS. urn:issn:0907-4449. 66, 125–132

77. Diederichs, K. (2006) Some aspects of quantitative analysis and correction of radiation damage. Acta Crystallogr D Biol Crystallogr. 62, 96–101

78. Liebschner, D., Afonine, P. v., Baker, M. L., Bunkoczi, G., Chen, V. B., Croll, T. I., Hintze, B., Hung, L. W., Jain, S., McCoy, A. J., Moriarty, N. W., Oeffner, R. D., Poon, B. K., Prisant, M. G., Read, R. J., Richardson, J. S., Richardson, D. C., Sammito, M. D., Sobolev, O. v., Stockwell, D. H., Terwilliger, T. C., Urzhumtsev, A. G., Videau, L. L., Williams, C. J., and Adams, P. D. (2019) Macromolecular structure determination using X-rays, neutrons and electrons: recent developments in Phenix. urn:issn:2059-7983. 75, 861–877

79. Emsley, P., Lohkamp, B., Scott, W. G., and Cowtan, K. (2010) Features and development of Coot. Acta Crystallogr D Biol Crystallogr. 66, 486–501

80. Afonine, P. v., Grosse-Kunstleve, R. W., Echols, N., Headd, J. J., Moriarty, N. W., Mustyakimov, M., Terwilliger, T. C., Urzhumtsev, A., Zwart, P. H., and Adams, P. D. (2012) Towards automated crystallographic structure refinement with phenix.refine. Acta Crystallogr D Biol Crystallogr. 68, 352–367

81. Chen, V. B., Arendall, W. B., Headd, J. J., Keedy, D. A., Immormino, R. M., Kapral, G. J., Murray, L. W., Richardson, J. S., and Richardson, D. C. (2010) MolProbity: all-atom structure validation for macromolecular crystallography. Acta Crystallogr D Biol Crystallogr. 66, 12–21

82. Grosse-Kunstleve, R. W., and Adams, P. D. (2003) Substructure search procedures for macromolecular structures. Acta Crystallogr D Biol Crystallogr. 59, 1966–1973

83. Terwilliger, T. C., Adams, P. D., Read, R. J., McCoy, A. J., Moriarty, N. W., Grosse-Kunstleve, R. W., Afonine, P. v., Zwart, P. H., and Hung, L. W. (2009) Decision-making in structure solution using Bayesian estimates of map quality: the PHENIX AutoSol wizard. Acta Crystallogr D Biol Crystallogr. 65, 582–601

84. Niesen, F. H., Berglund, H., and Vedadi, M. (2007) The use of differential scanning fluorimetry to detect ligand interactions that promote protein stability. Nat Protoc. 2, 2212–2221

85. Sassa, S. (1976) Sequential induction of heme pathway enzymes during erythroid differentiation of mouse Friend leukemia virus-infected cells. Journal of Experimental Medicine. 143, 305–315

86. Sinclair, P. R., Gorman, N., and Jacobs, J. M. (1999) Measurement of Heme Concentration. Curr Protoc Toxicol. 00, 8.3.1–8.3.7

