## Supplementary material for "Aim18p and Aim46p are CHI-domain-containing mitochondrial hemoproteins in *Saccharomyces cerevisiae*": Table S3

**Table S3. Data collection and refinement statistics.**

|  | **Aim18p (8ew8) (HREM)** | **Aim46p (8ew9)** |
| --- | --- | --- |
| **Wavelength (Å)** | 0.9644 | 0.9644 |
| **Resolution range (Å)** | 48.7–2.15 (2.227–2.15) | 40.06–2.0 (2.072–2.0) |
| **Space group** | P 3_2_ 2 1 | P 1 2_1_ 1 |
| **Unit cell (Å,** **°)** | 127.39 127.39 54.27  90 90 120 | 39.87 71.35 40.22  90 95.165 90 |
| **Total reflections** | 510482 (27578) | 102948 (10179) |
| **Unique reflections** | 27153 (2241) | 15174 (1471) |
| **Multiplicity** | 18.8 (12.3) | 6.8 (6.9) |
| **Completeness (%)** | 97.59 (81.11) | 99.61 (98.92) |
| **Mean I/sigma(I)** | 21.80 (2.38) | 11.15 (1.36) |
| **Wilson B-factor (Å^2^)** | 40.56 | 30.63 |
| **R-merge** | 0.09783 (0.9956) | 0.1234 (1.159) |
| **R-meas** | 0.1005 (1.039) | 0.1337 (1.253) |
| **R-pim** | 0.02266 (0.2836) | 0.05099 (0.4731) |
| **CC1/2** | 0.999 (0.702) | 0.998 (0.718) |
| **CC*** | 1 (0.908) | 0.999 (0.914) |
| **Reflections used in refinement** | 27146 (2241) | 15151 (1467) |
| **Reflections used for R-free** | 1988 (165) | 1521 (149) |
| **R-work** | 0.1781 (0.2409) | 0.1673 (0.2832) |
| **R-free** | 0.2125 (0.2701) | 0.2347 (0.3628) |
| **CC(work)** | 0.964 (0.837) | 0.971 (0.843) |
| **CC(free)** | 0.954 (0.750) | 0.942 (0.774) |
| **Number of non-hydrogen atoms** | 2061 | 2013 |
| **macromolecules** | 1908 | 1878 |
| **ligands** | 30 | 14 |
| **solvent** | 123 | 125 |
| **Protein residues** | 239 | 241 |
| **RMS(bonds)** | 0.007 | 0.008 |
| **RMS(angles)** | 0.80 | 0.84 |
| **Ramachandran favored (%)** | 97.05 | 98.33 |
| **Ramachandran allowed (%)** | 2.95 | 1.26 |
| **Ramachandran outliers (%)** | 0.00 | 0.42 |
| **Rotamer outliers (%)** | 0.46 | 0.48 |
| **Clashscore** | 1.55 | 1.85 |
| **Average B-factor** | 51.38 | 41.56 |
| **macromolecules** | 50.62 | 41.62 |
| **ligands** | 104.35 | 41.70 |
| **solvent** | 50.33 | 40.67 |
| **Number of TLS groups** | 6 | 7 |

Statistics for the highest-resolution shell are shown in parentheses.
